# Serial Imaging of Tumour and microEnvironment (SITE) platform for live-cell *ex vivo* modelling of primary and metastatic cancer dynamics

**DOI:** 10.1101/2025.09.08.674915

**Authors:** Vaibhav Murthy, Rawan Makkawi, Francis Anderson, Carol Halsey, Elise Manalo-Hall, Sanjay Srikanth, Gabriella Tangkilisan, Ting Zheng, James McGann, Marc R. Birtwistle, Jeremy Copperman, Alexander E. Davies

## Abstract

Understanding cancer initiation and progression is extremely challenging, in part due to experimental limitations in measuring and interpreting key signalling and tumour-microenvironment (TME) interactions that determine changes in cell and tissue behaviours over time. Here we developed SITE (Serial Imaging of Tumour and microEnvironment), a spatially-and temporally integrated, modular, *ex vivo* platform enabling quantitative analysis of TME interaction dynamics, signalling, and cell fate at single cell and tissue scales. Applied to modelling primary and lung metastatic breast cancer, SITE revealed tissue-specific TME interactions and ERK signalling patterns linked to distinct single-cell behaviours. We found that the earliest steps in tumour establishment and metastatic seeding involved active cell protrusion and the establishment of a multicellular niche interfacing tumour and host. Experimental and mathematical modelling showed that ERK signalling was co-influenced by these interactions, where cancer cluster formation increased signalling via the establishment of local signalling circuits. Disruption of these signalling circuits led to tissue-specific impacts on cancer intrinsic and TME interaction dynamics. Here, we modelled breast cancer as a test case, demonstrating the broad utility of SITE for quantitative exploration of TME interaction dynamics‒ closing a significant gap in experimental capabilities between *in vivo* models and *in vitro* systems.

## INTRODUCTION

Coordinated cell signalling and physical interactions within microenvironments are essential during development and maintenance, or regeneration, of tissue organization and function^1,2^. The mammary gland provides an elegant example of microenvironmental coordination across cellular and tissue scales. During post-natal development, hormonal cues induce expression of growth factors such as amphiregulin (AREG)^3–5^ and insulin-like growth factor 1 (IGF1)^6^. Establishment of these autocrine/paracrine signalling loops drives activation of key signalling pathways, e.g., ERK and AKT, that coordinate multicellular expansion and organized invasion to form the mature mammary gland structure^7^. Similarly, during pathologic insults such as wounding, signalling and physical interactions of epithelial, immune, and stromal cells, along with extracellular matrix (ECM) components drive coordinate restoration of normal tissue organization/function and appropriate response resolution^8–10^. In each case, these dynamic processes are unified by high fidelity perception and response to environmental signalling and interaction cues under tight regulatory control, dysregulation of which is a hallmark feature of disease, such as cancer initiation and progression^1,11^.

Primary and metastatic tumour microenvironments, or ‘host’ microenvironments, are structured yet heterogenous ecosystems comprised of cancer, resident, immune, and neuronal cells, vasculature, and ECM that coordinately influence cancer initiation and progression^1,12–15^. Signalling factors released into the tumour microenvironment, such as growth factors activating EGFR-Ras-ERK and other signalling pathways are often hijacked and/or amplified by tumour cells, leading to altered or inappropriate microenvironment perception-response dynamics and tissue organization. Simultaneously, tumour cells reprogram the host microenvironment, dysregulating ECM composition and structure, along with induction of host-cell states that can either suppress or favour tumour growth. It is now widely accepted that bidirectional tumour-host interactions dynamically evolve over time and their sum total, at the ecosystem or tissue-scale, regulate disease trajectory^16–18^. However, we still lack a deep understanding of how these interactions regulate trajectory, mechanistically.

A major challenge in understanding how tumour-host interactions precisely influence trajectory arises from a lack of tools that enable investigation of single cell-level behaviours, in the context of tissue, while also providing both spatial *and* temporal resolution. Single cell multi-omic sequencing, immunofluorescence imaging, and other approaches can identify diverse cell types and molecular states defined by gene expression^19,20^, chromatin state^21,22^, metabolites^23^, and protein abundance with single-cell resolution and even spatial context^24–27^, but fail to capture the dynamic nature of single-cells over time. Live-cell imaging techniques enable precise measurements of single-cell signalling and/or targeted gene or protein expression dynamics in 2D in-vitro culture^28–30^, and more recently in 3D in-vitro culture models^31–33^, but often lack important tissue-level context. Intravital imaging techniques have provided important insight into cell signalling and behaviour in living tissue environments, but experiments are difficult to perform, require very specialized equipment, and are low throughput^34–36^. An idealized system would retain structural and cellular features of the microenvironment, be experimentally facile, relatively high throughput, and provide high spatial and temporal resolution of tumour-host interactions underlying disease progression whilst integrating across cellular and tissue scales.

To address limitations of existing models, and questions underlying how tumour-host interactions influence tissue organization and steer disease trajectory, here we describe development and application of the Serial Imaging of Tumour and microEnvironment (SITE) platform. SITE enables live-cell ex vivo imaging of cancer cell dynamics within native tissue structures to provide tissue-specific context in organoid-tissue explant culture systems. SITE is coupled to fluorescent signalling biosensors and advanced live-cell imaging modalities, to simultaneously report cancer cell behaviours, physical interactions, and quantitative cell signalling activity in a single platform. These experimental models are complemented by a suite of computational approaches to extract single cell time series data with high precision and analyse cell fate and signalling trajectories in complex tissue microenvironments‒ resulting in end-to-end experimental and computational system, the SITE platform (**Figure 1**).

**Figure 1.**
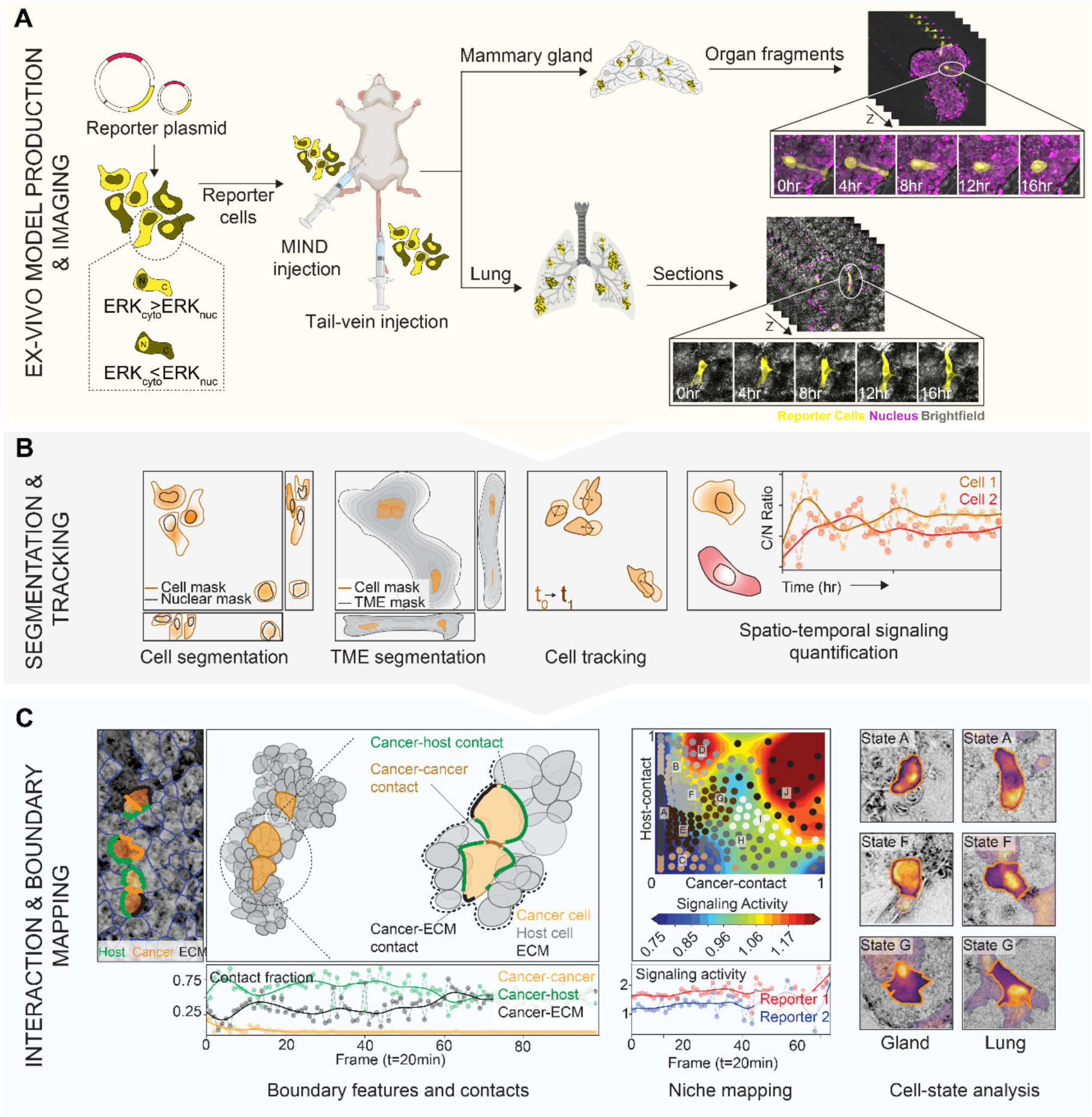
Overview of the Serial Imaging of Tumour and microEnvironment (SITE) platform. The SITE platform enables high-resolution, spatiotemporally resolved analysis of tumour–microenvironment interactions using ex vivo mammary gland (MammaSITE) and lung (LungSITE) tissue models. (**A**) Reporter breast cancer cells expressing fluorescent biosensors are introduced into the mammary duct via intraductal injection or lung via tail-vein injection, respectively. Post-injection, tissues are processed into organ fragments or precision-cut lung slices (PCLS) and imaged over time using 3D time-lapse microscopy to capture dynamic tumour behaviour. (**B**) Imaging data are processed using a computational pipeline for single-cell segmentation, tissue microenvironment classification, cell tracking, and quantification of ERK activity via cytoplasm-to-nucleus (C/N) ratio dynamics. (**C**) Spatiotemporal boundary mapping quantifies cancer cell contacts with neighbouring compartments (host cells, ECM, or other cancer cells), enabling construction of a high-dimensional contact feature space. These contact features are clustered to define niche states, which are linked to signalling activity and motility patterns over time across both mammary gland and lung tissue contexts.

To demonstrate the utility of the SITE platform, we applied it to experimentally model basal-like breast cancer at primary and metastatic locations. To this end, we generated <u>Mamma</u>ry gland and <u>Lung</u>-specific <u>SITE</u> models, the ‘MammaSITE’ and ‘LungSITE’, to investigate tumour cell signalling and tumour-host interactions at primary and metastatic locations, respectively. ERK biosensor expressing basal-like breast cancer cells were used to track cell signalling through this key pathway across tissue-specific contexts over time. We then explored the early steps of tumour formation in the mammary gland and dissemination to the lung. We monitored both signalling and tumour-host interaction dynamics simultaneously in these models. We found that the cancer cells initiated spatiotemporally coupled ERK signalling circuits, modulated directed protrusive motility to drive cancer cluster formation and extend the tumour-host interface and established distinct multicellular cancer-cancer and cancer-host local contact environments, or contact niches. Mathematical models derived from in-tissue single-cell ERK signalling measurements revealed that ERK signalling dynamics are tightly coupled to the signalling activity of adjacent cancer cell neighbours, consistent with the establishment of collective cell communities underlying tumour cluster organization, with implications for adaptation, survival, and growth decisions during the progression from primary to distant locations. The SITE platform and methods described here were successfully applied to experimentally model breast cancer progression, while specifically interrogating ERK signalling and tumour-host interaction dynamics. However, the tissue models, experimental, and computational approaches developed are applicable to a wide variety of cancer and tissue types amenable to *ex vivo* culture and could easily be paired with other existing or novel reporters to simultaneously capture and extract diverse signalling, tumour-host interaction, and/or cell fate dynamics.

## RESULTS

### A Serial Imaging of Tumour and microEnvironment (SITE) platform for multiscale live imaging of tumour-host dynamics

We designed the Serial Imaging of Tumour and microEnvironment (SITE) platform to integrate *ex vivo* organoid and tissue culture methods with fluorescent biosensor technologies, advanced live-cell imaging and computation into a single platform capable of simultaneously measuring and tracking cell signalling and tumour-host interaction dynamics at single cell and tissue scales (**Figure 1A-C**). The experimental workflow began with construction of cell lines co-expressing a biosensor of interest, followed by inoculation into the target organ to generate organ-specific tumour-host tissue models, in this case the mammary gland and lung (**Figure 1A**), MammaSITE and LungSITE, respectively. For MammaSITE, previous organoid derivation methods^37–40^ were modified to generate large organ fragments containing biosensor-expressing cancer cells with a target size ranging from ∼100 to 1000 µm (see Materials and Methods). For LungSITE, precision cut lung slice methods^41^ were used to generate 300 µm sections containing experimental metastasis of biosensor expressing cancer cells. Live-cell imaging data from MammaSITE and LungSITE were then paired with cellpose^42^ and ilastik-based software for segmentation^43^, tracking, and biosensor data quantification (**Figure 1B**), followed by application of computational methods to investigate complex single cell and tissue-scale relationships (**Figure 1C**), resulting in a complete experiment to analysis workflow.

### Establishment and validation of the SITE experimental platform to model primary and metastatic disease *ex vivo*

To establish and validate the SITE platform, we applied it to modelling intracellular signalling and tumour-host interaction dynamics in breast cancer. We chose the HMT-3522 T4-2 basal-like breast cancer cell line for our studies. T4-2 cells are adapted to the mammary gland *in vivo*, exhibiting rapid growth and local invasion within the primary site where they are significantly influenced by microenvironmental context via autocrine/paracrine EGFR-Ras-ERK signalling, most notably through EGFR ligands such as amphiregulin (AREG)^44,45^. Conversely, T4-2 cells are weakly metastatic undergoing significant attrition and often induction of dormancy following dissemination^46^. Thus, we reasoned T4-2 cells would provide a suitable model for understanding differential signalling and tumour-host interaction dynamics within the growth-promoting, adapted, environment of the mammary gland in contrast with the growth restricting, non-adapted, environment of the lung^45–47^.

To simultaneously visualize signalling dynamics, cell borders, and overall morphology of T4-2 cells, we engineered them to express an ERK kinase translocation reporter (ERK-KTR-mVenus) (**Figure 2A**), as previously described^48,49^. ERK-KTR biosensors are quantified by taking the ratio of cytoplasmic to nuclear (C/N Ratio) fluorescence intensities, with nuclear exclusion indicating increased ERK activity^48^. Biosensor expressing T4-2 cells were functionally validated by monitoring reporter activity under ERK activating and inhibiting treatments *in vitro* (**Figure 2A-B).** At both maximal and minimal reporter activation states cytoplasmic fluorescence remained detectable (**Figure 2A-B)** suggesting ERK-KTR may secondarily provide an accurate delineation of cell boundaries in tissue. To validate that ERK-KTR provided adequate resolution of cytoplasmic and cell membrane boundaries, we counterstained reporter expressing T4-2 cells with GAPDH (cytoplasm) and β-catenin (membrane-enriched). Regardless of baseline sensor activity, addition of ERK stimulating growth factors (EGF), or signalling inhibitors such as Erlotinib, we found that ERK-KTR provided accurate delineation cell boundaries (**Figure 2B**). Thus, we confirmed that ERK-KTR was suitable for detecting both signalling state and cell border/morphology, simultaneously.

**Figure 2.**
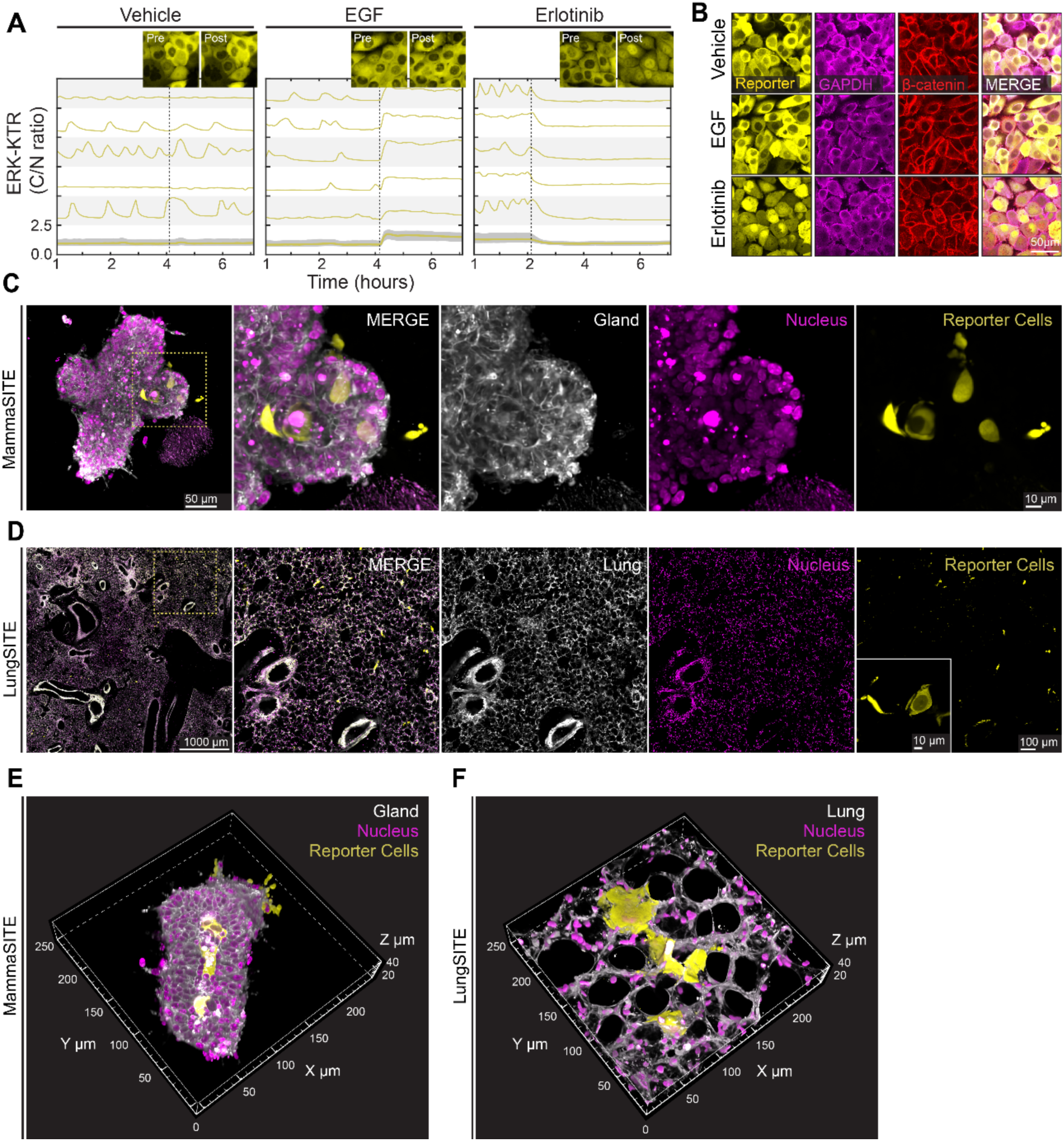
Functional validation and characterization of ERK biosensor-expressing breast cancer cells in *ex vivo* mammary gland and lung tissue model. (**A**) Measurements of ERK activity by cytoplasmic to nuclear ratio of ERK-KTR-mVenus reporter in T4-2 cells at baseline conditions (vehicle) or following treatment with EGF (100ng/mL) or EGFR inhibitor (Erlotinib, 2uM). In each panel, mean ERK activity is shown at bottom with the 25^th^–75^th^ interquartile range (IQR) depicted by shading from three biological replicates (N=3). Unshaded traces represent random single-cells and vertical lines indicate the addition of growth factor or inhibitor. Single-cell and mean ERK activity (n>2000 cells per condition) is plotted over 7-hour imaging (imaged every 6 minutes). (**B**) Representative 2-D images of ERK-KTR reporter distribution within T4-2 cells under baseline, EGF (100ng/mL) or EGFR inhibition (Erlotinib, 2µM). Cells were counterstained with cytoplasmic marker GAPDH (magenta), and membrane enriched β-catenin. Overlay images confirm that the reporter accurately delineates both the cytoplasmic compartment and cell boundaries. Scale bar=50µm, results shown from three technical replicates. (**C**) Representative maximum intensity projection of (left) whole mammary gland fragment showing successful engraftment of ERK-KTR reporter T4-2 cells (yellow) following mammary intraductal injection (MIND), and (right) zoomed in section of fragment highlighted in callout box. Scale bar= 50µm (left), 10µm (right). (**D**) Representative maximum intensity projection image of (left) PCLS, and (right) zoomed in section of white box showing successful engraftment of ERK-KTR reporter T4-2 cells (yellow) following tail vein injection to model metastatic dissemination to the lung. Inset shows high resolution image of reporter cancer cells. Scale bar=1000µm (left), 100µm (right), 10µm (inset). (**E**) 3-D rendering of mammary gland fragments showing ERK-KTR reporter cells (yellow) integrated into the gland lumen and interacting and invading through the luminal epithelium to the surrounding matrix. (**F**) 3-D rendering of PCLS illustrates distribution and integration of ERK-KTR reporter cells (yellow) along the host alveolar structure as shown by elongated morphology.

Validated biosensor expressing cells were injected into NOD SCID IL2R gamma null mice using the mammary intraductal (MIND)^50^, or lateral tail vein methods, to model primary site origination or metastatic dissemination, respectively (see diagram **Figure 1A**). Following injection, animals were humanely euthanized and mammary gland and lung tissues harvested. MIND injected mammary tissues were processed to generate organ fragments ranging from ∼100-1000 µM in size, aiming to preserve cellular composition and structural complexity of the gland. Lungs were prepared as 300 µm thick precision-cut lung slices (PCLS) to maintain native tissue architecture and cellular composition^41,51^ (**Figure 2C-D**). We observed that biosensor expressing T4-2 cells in the mammary gland readily integrated with the host tissue structure occupying the gland lumen and interacting with, or invading through, the luminal epithelium into the surrounding matrix (**Figure 2E**). In the lung, T4-2 cells formed complex interactions along host epithelial surfaces, often elongating over the alveolar walls, which appeared to guide cancer cell morphology (**Figure 2F**). Tracking of T4-2 cell viability as a function of reporter fluorescence, cellular morphology, and abundance in the MammaSITE and LungSITE revealed differential rates of tumour cell persistence and growth within these tissues. T4-2 cells underwent significant *ex vivo* expansion in mammary tissues, whereas in the lung they underwent marked attrition by day 7 post-injection (**Supplementary Figure 1A)**. These data are consistent with previously reported *in vivo* growth characteristics of the T4-2 cell line in the mammary gland and lung^45^. To confirm the modularity of SITE approaches to other cell lines and tumour types we generated MDA-MB-231 triple-negative breast cancer reporter lines to express ERK-KTR, as described for T4-2 cells, and OS-17 osteosarcoma cells^52^ to express either ERK-KTR or co-express ERK-KTR, an AKT sensor^53^, and a histone marker and generate ex vivo tissue bearing these cells (**Supplementary Figure 1B-D**). As expected, MDA-MB-231 ERK-KTR cells were readily detectable and remained viable in mammary and lung tissue for 7-days when collected immediately after injection (**Supplementary Figure 1B**). We additionally grew MDA-MB-231 and OS-17 osteosarcoma cells in vivo for 14-days before lung harvest demonstrating that SITE can be performed at alternative time points (**Supplementary Figure 1C**). Further, OS-17 cells co-expressing multiple reporters in lung were harvested under these conditions demonstrating biosensor multiplexing capabilities (**Supplementary Figure 1D**). Finally, host tissue viability was verified by mitochondrial function and cell viability assays. In both mammary and lung tissues we found robust viability in host cell populations for up to the maximum time point evaluated, 7 days, *ex vivo* (**Supplementary Figure 1E)**. Together, these data indicate that SITE models and modular and can be maintained with high viability *ex vivo* to recapitulate expected *in vivo* tumour cell growth-persistence behaviours while maintained in culture (**Supplementary Figure 1E**).

### Development and validation of the SITE image analysis pipeline

To support computational pipeline development, MammaSITE and LungSITE live-cell imaging datasets were collected for each tissue. Immediately post-injection, images were captured every 20 minutes for a period of 24 hours. Media conditions included control (no growth factor or inhibitor added), or MEK inhibitor (MEKi, PD0325901, 500nM) to suppress ERK activity. Imaging data revealed a wide variety of tissue architectures, cancer cell cluster sizes, tumour-host interactions, and time-varying signalling information (**Figure 3A**). To analyse these data, we first extracted imaging subvolumes from larger datasets to focus on regions of interest (ROIs), minimizing file size and overall processing time. Next, we applied deep learning-based segmentation approaches to delineate cytoplasmic and nuclear compartments using cellpose software^54^ that was manually trained to identify and segment these features in 3-dimensions (**Supplementary Figure 2A, Figure 3B**). To further delineate tissue boundaries and extracellular matrix (ECM) we applied a sparse labelling approach in a semi-supervised framework using the ilastik software package that was manually trained (**Figure 3B-D**)^43^. In this context, we defined ECM as any non-cellular contact interface. To ensure segmentation fidelity, a subset of images were manually verified and compared to the automated segmentation results, confirming delineation accuracy for cellular (mammary gland: 89.6%, lung: 85.7%, **Supplementary Figure 2A-B, Supplementary Table 2, Supplementary Table 3**) and microenvironmental components (percent pixel-level overlap—mammary gland: 86%, lung: 70%, **Supplementary Figure 2C, Supplementary Table 4**). Cytoplasmic segmentation masks were then linked through time using an optimal transport-based approach^55^ (**Supplementary Figure 2A, Supplementary Table 2)**. The result was an easily accessible data framework for subsequent investigation of features such as cell contact surfaces, cell-cell interactions, directional movement and motility, and ERK reporter dynamics obtained from SITE tissue models (see diagram **Figure 1B**).

**Figure 3.**
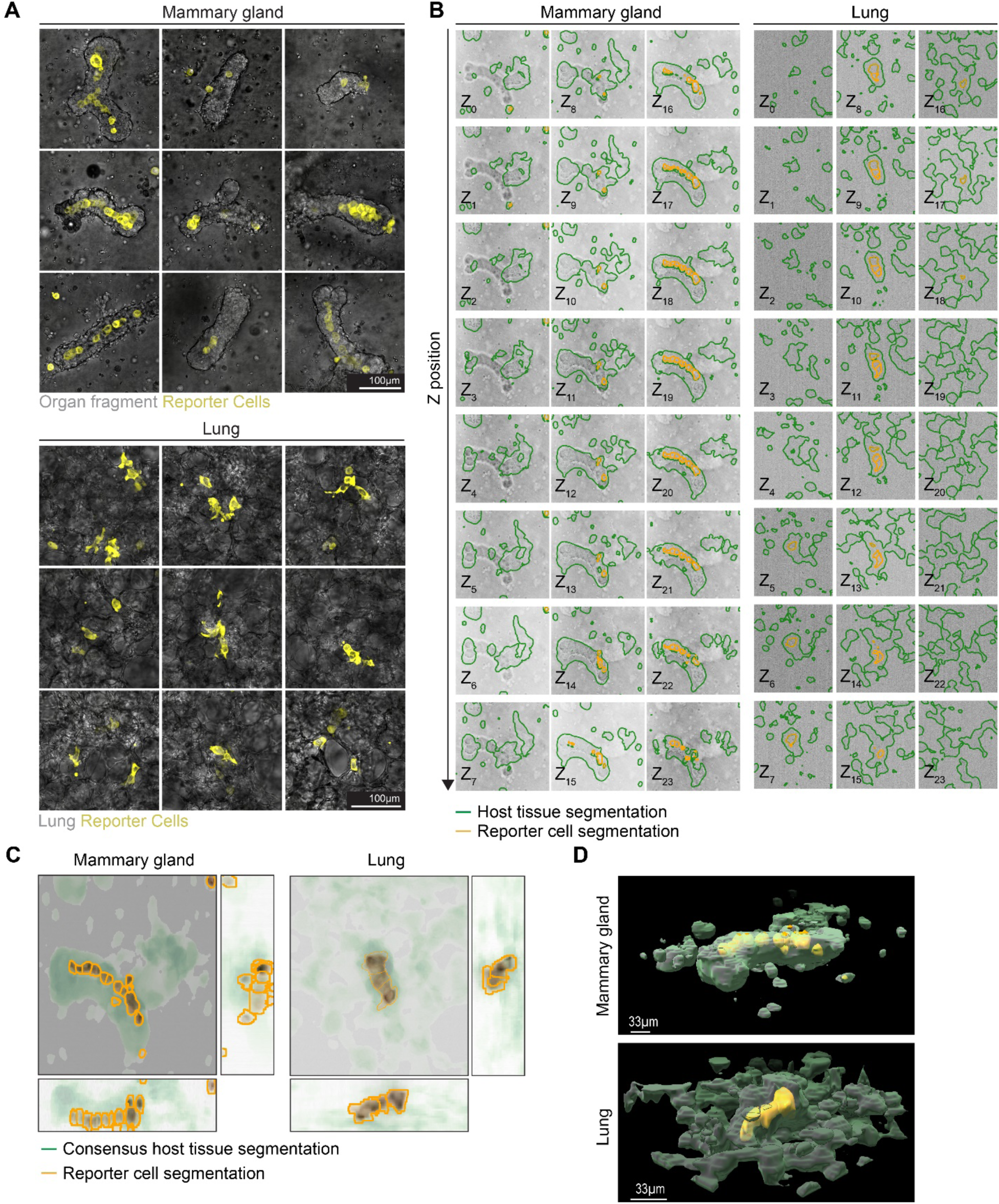
Automated segmentation of tumour-host interactions in ex-vivo mammary gland and lung models. (**A**) Representative maximum intensity projection images of ERK-KTR reporter T4-2 cells (yellow) within mammary organ fragments (grey, top) and PCLS (grey, bottom) captured immediately post-injection following tissue preparation. Images depict a range of tissue architectures and reporter cell cluster sizes, highlighting the versatility of the SITE platform for capturing diverse tumour-host microenvironments. Scale bar=100µm. (**B**) Individual Z-stack slices illustrating segmentation of reporter cells (orange outlines) and host tissue (green outlines) in mammary gland organ fragments (left) and PCLS (right). Segmentation was performed using machine learning-based approaches (Cellpose for reporter cells, n>1000 training images; ilastik for host tissue, n>100 training images), enabling accurate delineation of cellular and microenvironmental compartments across multiple focal planes. (**C**) Overlay of consensus segmentations (described in Methods) for reporter cells (orange) and host tissue (green) in representative regions of interest from mammary gland organ fragment (left) and PCLS (right). (**D**) 3-D rendering of the composite individual slice segmentations of reporter cells and host tissue for mammary gland organ fragments (top) and PCLS (bottom). The rendering integrates segmented data across serial Z-stacks providing a comprehensive view of cellular and tissue architecture. Scale bar= 33µm

### Protrusive cell motility drives formation of tissue-specific contact niches

With the experimental and computational frameworks in place, we first used SITE to understand how microenvironmental context affects cancer cell organization and behaviour in mammary gland and lung models by live-cell imaging over 24 hours. In the mammary gland, we observed that cancer cells exhibited a tendency to move from their initial positions through the formation of protrusions, often while migrating toward other cancer cells (**Figure 4A**, top). Similarly, in the lung, individual cancer cells often formed protrusions towards cancer cells, with protrusions directed along alveolar structures (**Figure 4A**, bottom). To better quantify this observation, and the directionality of cancer cell motility, we calculated the directed displacement of cells over time and distance, as diagrammed in **Figure 4B** (top). Relative displacement was calculated as the centroid displacement of an individual cancer cell to neighbouring cancer cell(s) separated by a distance r and time t, hereafter referred to as relative motility. As such, we were able to evaluate the net motility of a cancer cell toward or away from cancer neighbours (positive and negative displacement, respectively) in space and time. Using this approach, we observed relative motility towards other cancer cells in both the gland and lung microenvironments at cell separations between ∼20-60 µm over time periods exceeding 5 hours (**Figure 4B**, bottom, **Supplementary Figure 3B-E**). These data provided evidence of self-organization over extended time periods during early tumour formation. At distances of several cell lengths (>60 µm), directed motility was diminished, suggesting largely local chemotactic behaviour via secreted mediators in this tissue setting. This range was relatively unaltered by MEK inhibition (**Supplementary Figure 3A-3D**), suggesting a level of independence between ERK activity and cancer-directed relative motility. At separations less than 10 µm, we observed a net negative displacement suggesting a repulsive-like interaction driving relative motility of a cancer cell away from its neighbour, consistent with engagement of cell-cell spacing and tissue-patterning mechanisms.

**Figure 4.**
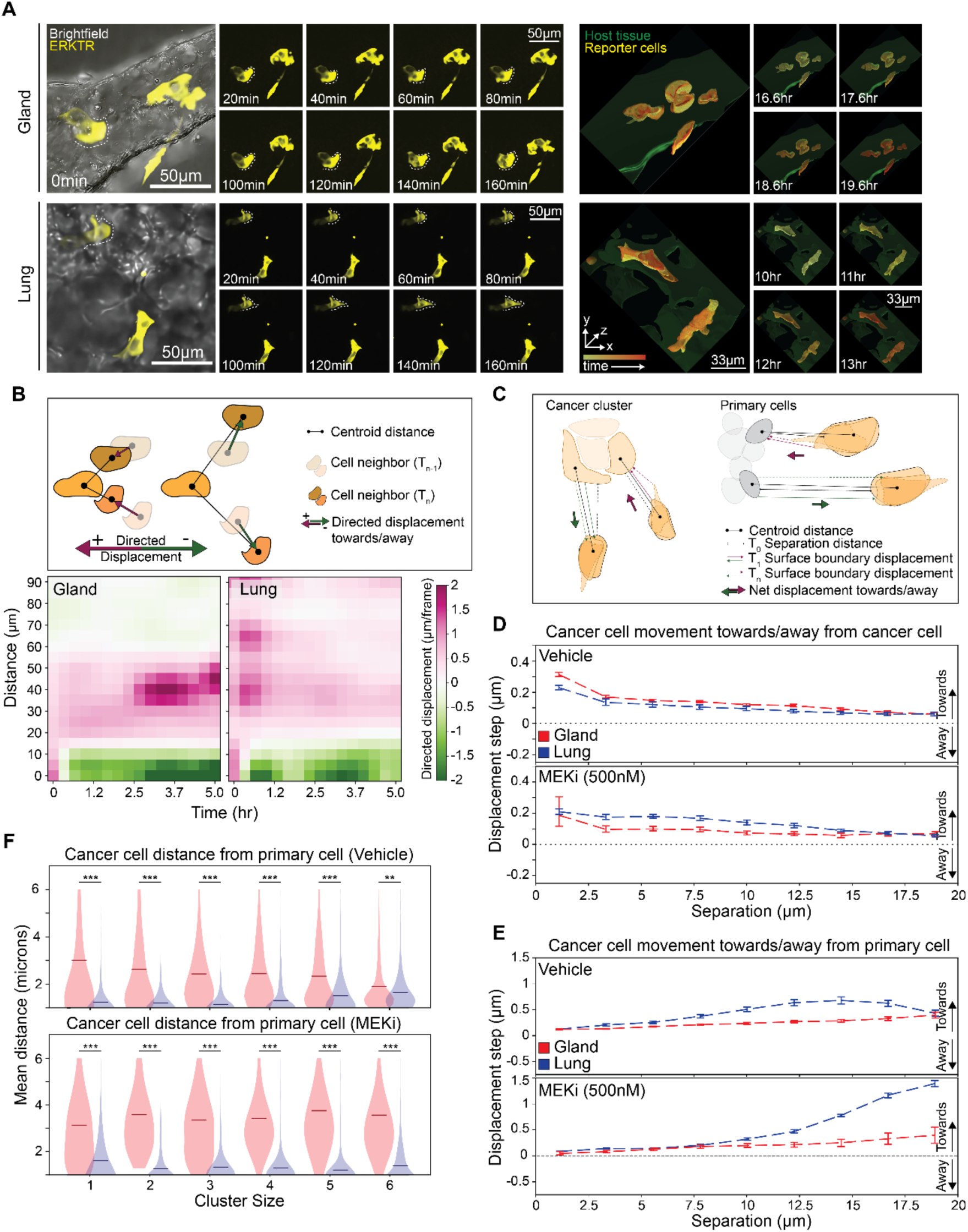
Protrusive cell motility drives formation of tissue-specific contact niches. (**A**) Representative time-lapse maximum intensity projections (left) of ERK-KTR reporter T4-2 cells (yellow) in mammary gland organ fragments (top, grey) and PCLS (bottom, grey), acquired at 40X magnification every 20 minutes over 24 hours, with ERK-KTR images from initial timepoints and boundary undergoing protrusive motion highlighted (white dashed line). Scale bar=50µm. (Right) 3-D rendering of reporter masks and host tissue masks showing cell movement and motility at represented timepoints. Merged renderings overlay individual-coloured masks to show maximal displacement over 4-hour window. Scale bar=33µm. (**B**) Schematic illustration (top) of centroid displacement calculation of individual cancer cells over time relative to their nearest cancer cell neighbour. Pink and green arrows designate net positive and net negative centroid displacement of cell towards or away from other cancer neighbours. Quantitative heatmaps (bottom) show aggregated net cell movement as a function of initial separation distance. (**C**) Localized boundary displacement schematic for each cancer cell relative to nearest cancer or host cell neighbour. Boundary displacement to the next frame was projected onto the direction of the nearest neighbour to quantify directed motility. (**D**) Cancer cell boundary displacement towards or away from nearest cancer cell neighbour in vehicle (top) or MEK inhibitor (PD0325901, 500nM) treated conditions in mammary organ fragments (red) and PCLS (blue). Graphs are plotted as mean (dashed lines) and bootstrapped 95% CI over biological replicated. (**E**) Cancer cell boundary displacement towards or away from nearest host cell neighbour in vehicle (top) or MEK inhibitor treated conditions in mammary organ fragments (red) and PCLS (blue). Graphs are plotted as mean (dashed lines) and bootstrapped 95% CI over biological replicated. (**F**) Violin plots of cancer–host centroid distances, stratified by cluster size in mammary organ fragments (red) and PCLS (blue) in vehicle (top) and MEK inhibitor treated (bottom) conditions. Solid bars represent mean of all plotted cells within the cluster size classification (see Methods). Significance was determined by means of Welch’s t-test (***-P≤0.0001, **-P≤0.001) on individual ROIs. A summary of pairwise statistics can be found in Table 5.

We next investigated motility, cancer-cancer, cancer-host interaction relationships in more granular detail (**Figure 4C-E**). As in the previous example, we calculated displacements over time; however, we applied this calculation to cancer cells relative to each other *and* relative to adjacent host cells. Additionally, we calculated displacements directly at the surface boundary for each point on the cell boundary relative to its nearest neighbouring cancer cell or host cell, rather than the coarser centroid level. This allowed us to extract quantitative cell-cell interaction dynamics e.g., contact of cell protrusions at the membrane deformation level, complex features that were not captured in our previous analysis (see diagram **Figure 4C**). We observed an average positive displacement at the cancer cell boundary towards the nearest neighbouring cancer cell over a range of ∼20 µM, with cancer-to-cancer cell protrusive motility increased in the gland compared to the lung under control conditions (**Figure 4D**, top). Simultaneously, we observed average positive displacement at the cancer cell boundary towards the nearest host cell extending over ∼20 µm (**Figure 4E**, top). This attractive displacement was increased in the lung compared to the gland, peaking at a separation of ∼15 µm, indicating increased attractive cancer-host interactions in the lung microenvironment. Blocking ERK signalling with MEK inhibitor, reduced cancer-to-cancer protrusive motility in the gland but not in the lung (**Figure 4D**, bottom). In contrast, MEK inhibition abrogated cancer-to-host protrusive motility in the gland but significantly increased this protrusive motility in the lung (**Figure 4E**, bottom). Together, these data suggest that the effects of silencing cancer autonomous signalling programs are tissue-dependent between primary and metastatic sites, with ERK activity coordinating the spatial tumour architecture through a balance between niche-specific cancer-cancer and cancer-host communication and interaction.

Given these data, we reasoned that the observed cell boundary displacement properties should play a role in establishing the architectural properties (i.e., clustering) of the early tumour formation microenvironment. To evaluate this, we measured the relative distributions of cancer and host cells in the MammaSITE and LungSITE models as a function of cancer cluster size. As expected, in both mammary gland and lung, the mean distance between the cell boundary of a cancer cell to their nearest neighbouring cancer cell decreased as a function of increasing cluster size (**Supplementary Figure 3E**). Once cancer-cancer direct contact was established for 2 cells, the mean distance to other cancer cells was largely similar with increasing cluster size, indicating the maintenance of balanced cancer-cancer and cancer-host contacts as the clusters increased in size. However, clusters of cancer cells in the lung established increased contacts with host cells relative to the gland when clusters were comprised of 4 or less cells (**Figure 4F**, top). This was because cancer cells in the lung tended to form elongated structures resulting in significant host-contact interfaces that appeared to be guided by the lung structure (**Figure 4A**, bottom). In contrast, in the gland, cancers cells on average were furthest from host cells when cancer clusters had only 1 or 2 cells corresponding to a subpopulation of cells located towards the periphery of the organ fragment and the ECM (**Figure 4A**, top). MEK inhibition accentuated these tissue-specific differences in multicellular cancer clusters of 5 or more cells, observed as increased distance to host cells in the gland, and decreased distance to host cells in the lung. Altogether, these data indicate that small cancer clusters adopt distinct niches in the lung and the gland, and that differential protrusive motility towards host cells in these organs contributes to the formation of tissue-specific niche structures and invasive behaviours.

### ERK signalling is spatially and temporally coordinated by intercellular communication between cancer cells

The protrusive motility we observed demonstrates how cancer cells adaptively respond to and shape their microenvironment, establishing early contact niches that are distinct between the gland and the lung. We found that these early contact niches are dynamic suggesting they may be modulated by intracellular signalling states across local neighbourhoods of cancer cells. Since ERK signalling in T4-2 cells is known to be driven in part by autocrine/paracrine mechanisms, whereby secreted AREG ligands engage EGFR receptors on adjacent cells, initiating a positive feedback loop (diagram **Figure 5A**),^56^ we next sought to understand how ERK activity in one cell could drive signalling in a neighbouring cell via soluble mechanisms. To do so, we first quantified ERK activity patterns as a function of intercellular distance, finding that high ERK activity in one cancer cell was highly correlated with ERK activity in neighbouring cancer cells, in both the lung and the gland microenvironments (**Figure 5B-C, Supplementary Figure 4A-B**). This finding was consistent with previous reports of spatial ERK signalling coordination via soluble ligand inducing activity waves within cellular communities during wound healing and tissue development^57–59^. We observed signalling coordination at distances of >60µm that was maintained over extended durations (>5 hours). ERK-mediated long-distance signalling was dampened upon MEKi treatment independent of the model system (**Figure 5B, Supplementary Figure 4A-B**). Together, we interpreted these data as suggesting a role for ERK-induced soluble factor secretion itself mediating these long-range communication events.

**Figure 5.**
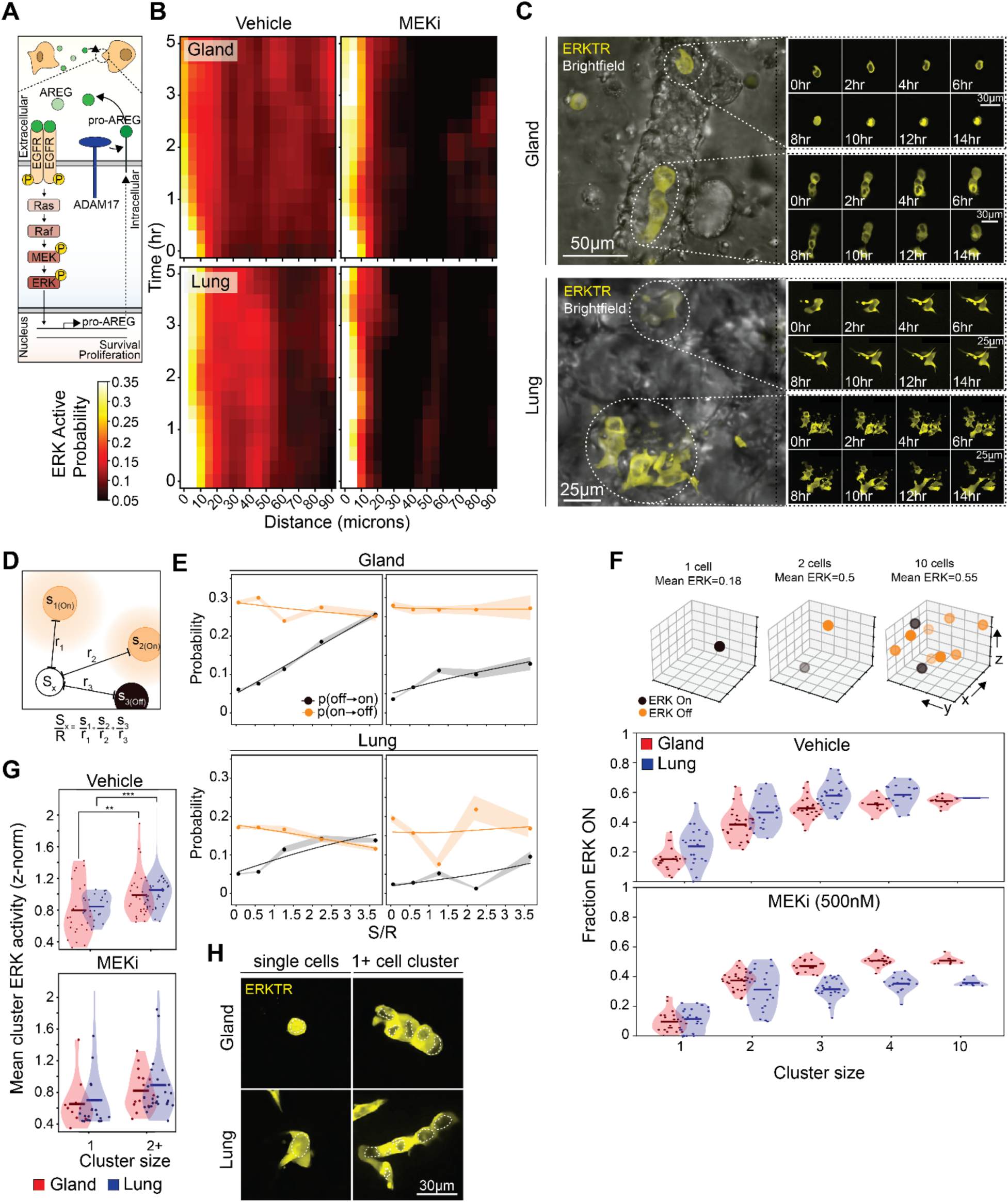
ERK signalling is spatiotemporally coordinated by intercellular communication and modulated by local ERK-active cell density. (**A**) Schematic illustration of autocrine/paracrine ERK signalling, depicting amphiregulin (AREG) secretion from HMT-3522 T4-2 cells activate EGFR on neighbouring cells, initiating a positive ligand-mediated feedback loop. (**B**) Heatmaps of spatiotemporal ERK activation in vehicle vs. MEK inhibitor (PD0325901, 500nM) treated conditions. Graphs represent ERK activation probability as a function of time and distance from neighbouring ERK_ON_ cell. (**C**) Representative maximum intensity projection images of time-lapse imaging of ERK-KTR reporter T4-2 cell (yellow) signalling dynamics in mammary gland organ fragments (grey, top; Scale bar=50µm) and PCLS (grey, bottom; Scale bar= 25µm). Callout panel shows maximum intensity projections of ERKTR reporter activity of single cells and clusters tracked over time in corresponding tissue models. Scale bar=30µm (gland), 25µm (lung). (**D**) Schematic of signalling metric S/R quantification based on inverse distance-weighted ERK-on neighbours. ERK activation rates increase with higher local density of ERK-on neighbours. ERK signalling state in target cell (white) calculated based of sum of number of ERK_ON_ (orange) and ERK_OFF_ (black) cells and their distance (r) from target cell. (**E**) Graphs of probability of ERK activation (OFF→ON, orange) and deactivation (ON→OFF, black) as a function of local signalling density (S/R, signal-to-radius) in vehicle vs MEKi treated conditions in mammary organ fragments (top) and PCLS (bottom). Probability plotted as mean (circles) and bootstrapped 95% CI over biological replicated (shaded region) with solid lines as guides to the eye. (**F**) Diagram of (top) Markov modelling of ERK activation rates as a function of local neighbour signalling density and violin plots (bottom) of predicted ERK_ON_ fraction of cells stratified by cluster size in vehicle vs. MEKi treated conditions in mammary organ fragments (red) and PCLS (blue). (**G**) Violin plots of z-normalized mean ERK activity in single cells vs. clusters in vehicle vs. MEKi treated conditions in mammary organ fragments (red) and PCLS (blue). Solid bars represent the mean, and points represent individual cells/clusters tracked and significance was determined by means of Welch’s t-test (***-P≤0.0001, **-P≤0.001). (**H**) Representative maximum intensity projection images of small/single and large reporter cell clusters in mammary organ fragments and PCLS. White dotted markings indicate nuclei showing ERK reporter localization (cytoplasmic vs. nuclear). Scale bar=30µm.

To further understand how signalling dynamics are spatiotemporally regulated based on neighbouring cancer cell context, we developed a spatial signalling metric (S/R; see methods for equations), where S represents the ERK on/off state and R represents the distance to nearby cancer-cells, that captures the influence of ERK-active neighbours (diagram **Figure 5D**). By following single cell timeseries of ERK activation states, we derived ERK activation state transition probabilities, in the context of the S/R spatial signalling metric. In untreated conditions, ERK activation probability (ERK_OFF_→ERK_ON_) increased with higher local signalling density in both gland and lung environments suggesting that ERK state changes were dependent on the sum of the number of ERK active signalling neighbours and their distance, where increasing the number of ERK_ON_ signalling cells increases the probability of an ERK state transition change. As expected, the addition of MEK inhibitor in both the gland and the lung reduced ERK_ON_ rates and significantly dampened the increase of ERK_ON_ rate with local signalling density (**Figure 5E**).

Since both cancer-exogenous and cancer-autonomous signal variation influence cancer cell ERK activation dynamics, we applied a Markov model defined by extracted ERK_ON_ and EKR_OFF_ rates, to isolate how cancer-autonomous paracrine signalling alone influences ERK activation patterns (**Figure 5F**). The model, via the local signalling density, incorporates spatial coupling of ligand signalling to capture quantitatively how ERK activity would increase as a function of cluster size, with tissue and treatment specific kinetics. To validate the model prediction of increased ERK activity in clusters of cancer cells compared to isolated cancer cells, we measured mean ERK activity across different cluster sizes and found that multicellular clusters indeed exhibited higher ERK activity than isolated cells (**Figure 5G-H**). This relationship was apparent in both gland and lung environment. Together, these results establish that ERK signalling is spatially coordinated through across tissue scales, with cancer cells likely modulating the signalling activity of their neighbours via ERK-mediated paracrine signalling, which carries implications for overall niche organization.

### Cell-cell contact-defined local niche states influence cancer cell motility and ERK signalling

Our cell motility analysis revealed how cancer cells actively shape tissue-specific contact niches through interactions with both other cancer cells and host cells. We found that these interactions vary between the gland and the lung and, in part, modulate ERK signalling activity in the corresponding tissue suggesting that cellular contact niches could produce both structural and functional outcomes. To systematically explore these relationships further, we analysed cancer cell behaviours as a function of their contact niche. We treated contact niches as unique microenvironments, asking how distinct ratios of cancer-cancer, cancer-host, and cancer-ECM contacts (**Figure 6A-B**) influence cancer cell states (behaviour and signalling) in distinct and measurable ways over time. To this end, we first applied the SITE computational pipeline (**Figure 3**) to extract tumour-microenvironment interactions by performing contact feature analysis to delineate cancer cell contacts with other cancer cells and host cells (**Figure 6A**, left). Each observed single-cell was mapped into a 2D contact space quantifying cancer-cancer and cancer-host contacts. Single-cell trajectory timeseries of these contact features informed a trajectory-based kinetic clustering, as previously described^60^, resulting in ten distinct “contact niche states” (represented as states *A-J,* **Figure 6A-B**) each defined by different contact relationships. Because presentation of contact niche states relationships is complex and often abstract, we generated ‘in tissue’ plots to better visualize these states and their spatial relationships to the host (**Figure 6C**). These plots delineated cell states and features by position within in the gland and lung (**Figure 6C**, representative single time point examples), capturing snapshots of dynamic states *A-J* from live-cell imaging data and were output in movie format to visualize evolution of contact niche state relationships over time (**Supplementary Figure 5A)**. We then calculated relationships between contact niche states and cell motility using frame-by-frame centroid displacement in microns and signalling activity calculated as the cytoplasmic to nuclear fluorescence ratio (C/N ratio) of the ERK reporter. To explore the link between contact niche state and cell behaviour, state centres were superimposed on heatmaps of motility or ERK reporter activity, in the space of cancer-host and cancer-cancer contact fractions (**Figure 6D**). Combining MammaSITE and LungSITE data together revealed common trends that increased cancer cell motility was associated with higher cancer-cancer contact fractions (states *H-J*) (**Figure 6D-E)**. States *H-J* also exhibited significantly elevated ERK activity; with the highest ERK activity level associated with cells in state *J* (**Figure 6E)** that simultaneously display high cancer-cancer *and* cancer-host contact fractions (**Figure 6D-E**), consistent with a paradigm of ERK-stimulating paracrine signalling converging on a privileged subpopulation of cells towards the tumour periphery.

**Figure 6.**
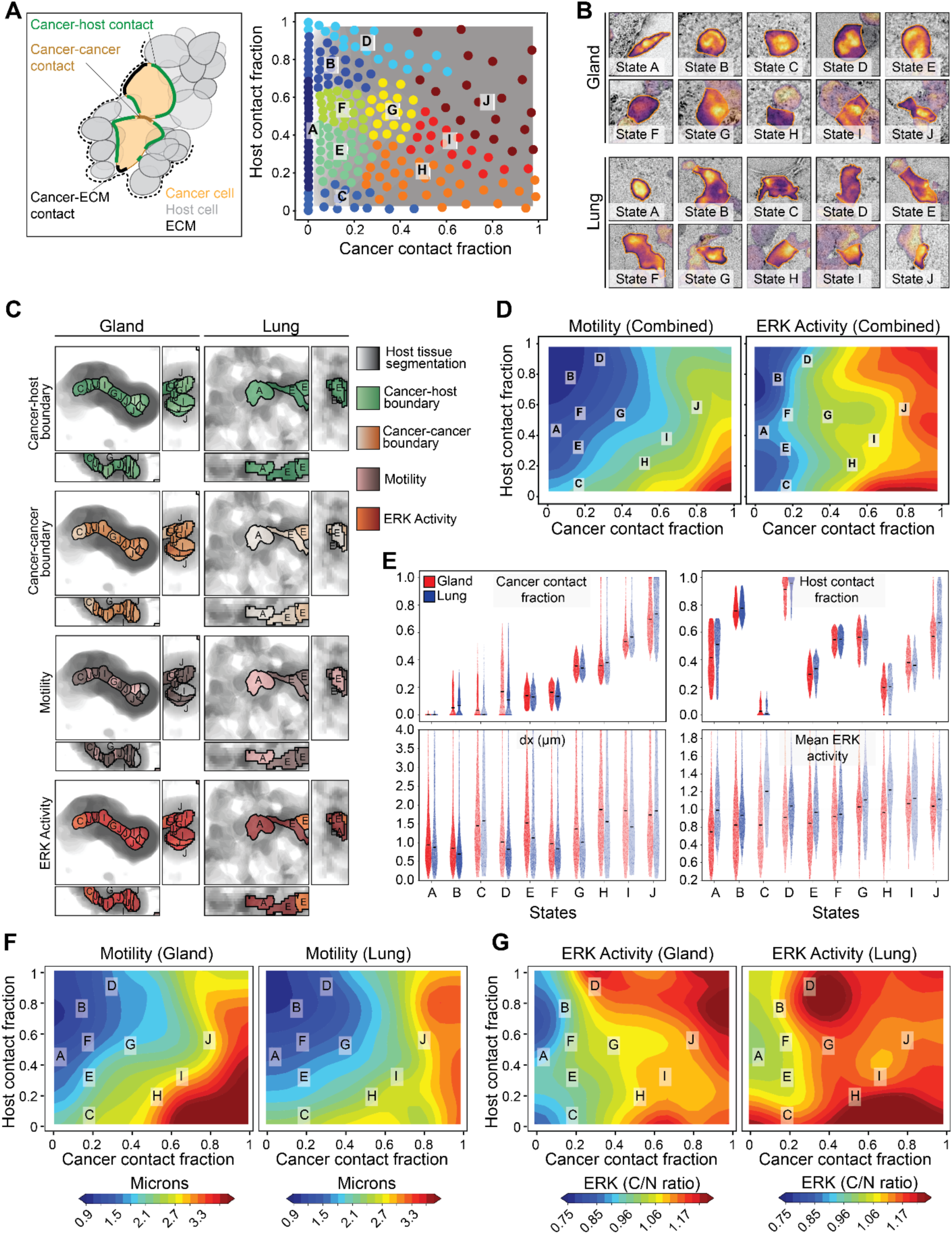
Tumour–microenvironment interaction features define contact niche states that correlate with cell motility and ERK signalling dynamics. (**A**) Diagram (left) of boundary features computed from 3D segmentation masks, classifying cancer cell contacts with neighbouring cancer cells (orange), host tissue cells (green), or ECM (grey). Contacts were defined by a nearest-neighbour distance ≤1.9 μm (see Methods). 2-D latent space (right) of ten discrete metastable contact niches (A-J) obtained from trajectory-informed kinetic clustering (k-means, n=220) of cancer-cancer (x-axis) and cancer-host (y-axis) in combined vehicle and MEKi (PD0325901, 500nM) mammary gland organ fragment and PCLS conditions. (**B**) Example single-cell images from each contact niche state (A–J) in the gland (top two rows) and lung (bottom two rows) microenvironments. Cells are outlined and coloured by ERK activity. (**C**) Representative spatial maps from the mammary gland organ fragments (top) and PCLS (bottom) models showing assignment of individual cancer cells to contact niche states based on boundary features. Columns display, from top to bottom: cancer-host boundary (green), cancer-cancer boundary (orange), motility (pink), and ERK activity (red). Each cell is color-coded and labelled according to its assigned contact niche state (A–J). (**D**) Motility (left) and ERK activity (right) map associated with contact niche states across all tissue models and conditions (vehicle and MEKi). States A/B and J exhibit the lowest and highest motility, respectively. States D and J-characterized by high contact with either cancer or host cells exhibit elevated ERK signalling. (**E**) Violin plots of (top-left) cancer contact surface fraction and (top-right) host surface contact fraction for each state represented in (**A**) for mammary organ fragments (red) and PCLS (blue). Cumulative centroid displacement plots (bottom-left) and mean ERK-KTR cytoplasm-to-nucleus (C/N) ratio (bottom-right) in each contact niche state. (**F**) Tissue-specific motility maps with superimposed contact niche states (A-J). (**G**) Tissue-specific ERK activity maps with superimposed contact niche states (A-J).

Finally, we performed a tissue specific analysis, splitting gland and lung data to obtain unique features within each, then plotting contact niche states as a function of cancer-cancer and cancer-host contact fractions and superimposed on heatmaps of motility (**Figure 6F, Supplementary Figure 5B)** and ERK signalling (**Figures 6G, Supplementary Figure 5C)**. As observed for the combined analysis, cancer cells exhibited the highest motility with a high cancer-cancer contact with variable cancer host-contact fractions (**Figure 6F**) indicating similarity in cancer cell motility behaviours in both gland and lung contexts. However, in the gland, high ERK activity trended toward cells with high cancer and host contact fractions (contact niche state *J*) (**Figure 6F**), whereas in the lung, high ERK activity was more widely distributed amongst the contact niche states (contact niche states *D, G, H-J*) (**Figure 6G**). Together these data highlight how different tissue microenvironments can similarly impact cancer cell motility, whilst having more divergent effects on cell signalling, demonstrating that ERK signalling can be at least partially decoupled from cell motility depending on environmental context.

## DISCUSSION

Here we described a novel integrated experimental and computational platform, the Serial Imaging of Tumour and microEnvironment (SITE) platform (**Figure 1**), coupling biosensors, *ex vivo* tissue models and advanced live-cell imaging with computation to track and interrogate dynamic tumour-host interactions. Using this platform, we demonstrated its ability to recreate features of the primary mammary and metastatic lung breast cancer microenvironments (MammaSITE and LungSITE, respectively), extracting and quantifying cell motility, cancer-cancer, cancer-host, and signalling interaction features over time and in 3-dimensions (**Figure 2**, **Figure 3**).

Using the MammaSITE and LungSITE models, we found that early tumour forming cells exhibited directional protrusive motility (**Figure 4A-E)** and establish distinct multicellular niches within the luminal space in mammary gland and inter-alveolar space in the lung (**Figure 2E-F**, **Figure 4F**). These niches were stratified based on the interactions cancer cells make with other cancer or host cells (**Figure 4A-E**, **Supplementary Figure 5**). A consequence of these niches was the establishment of ERK signalling circuits, that were themselves, spatiotemporally regulated by ERK signalling-induced secretion. In our models, this apparent cancer-cancer autocrine/paracrine signalling loop led to the observation of increased motility and ERK signalling with increased cancer contacts in both primary and metastatic environments (**Figure 6E-G**). In relation to host contact associated ERK signalling in cancer cells, we found distinct behaviour between the mammary gland and lung, which was likely to be driven by distinct and unresolved lung host cell types in our live imaging. A potential mechanism is growth factor secretion from stromal cells at the tumour-stromal boundary, which has emerged as a key driver of invasion^61^. Additionally, our findings point to intercellular paracrine communication between cancer cells to also play a crucial role in coordinating converging ERK signals at the tumour periphery.

The shared trajectory-based state-space framework revealed that while both motility and ERK signalling drive the emergence of dynamic contact niches (**Figure 6D)**, the structure of these niches is strongly influenced by the host tissue environment. Detailed analyses of motility and signalling demonstrated that the interplay between cell-intrinsic signalling and cell-extrinsic context is central to shaping cancer cell behaviours, resulting in distinct patterns between the mammary gland and the lung (**Figure 6F-G**). In both tissue models, cancer cells exhibited spatiotemporally coordinated ERK signalling activity, likely mediated by ERK-dependent secretion from neighbouring cells—a phenomenon disrupted by MEK inhibition (**Figure 5A-B**). Elevated cancer-cancer contact interfaces and increased ERK signalling consistently promoted the formation of multi-cellular clusters across both the mammary and lung microenvironments (**Figure 4B, 5G-H**), yet the outcomes of these interactions diverged depending on the tissue context. Specifically, cancer cells in the mammary gland favoured compact architectures rich in cancer-cancer contacts (**Figure 4D**), while the structural characteristics of the lung gave rise to elongated clusters with increased cancer-host contact interfaces (**Figure 4E**). MEK inhibition further accentuated these differences—reducing protrusive cancer-host motility in the gland, while paradoxically enhancing it in the lung (**Supplementary Figure 5B-D)**.

Collectively, these findings are summarized in **Figure 7** and highlight the context-dependence of ERK-driven behaviours and the remarkable plasticity that characterizes metastatic adaptation. Furthermore, they underscore the power of SITE to dissect and map in real-time how contact-and signalling-dependent mechanisms orchestrate niche formation in both primary and metastatic tissues and advance our understanding of tumour heterogeneity and microenvironmental influence in cancer progression.

**Figure 7.**
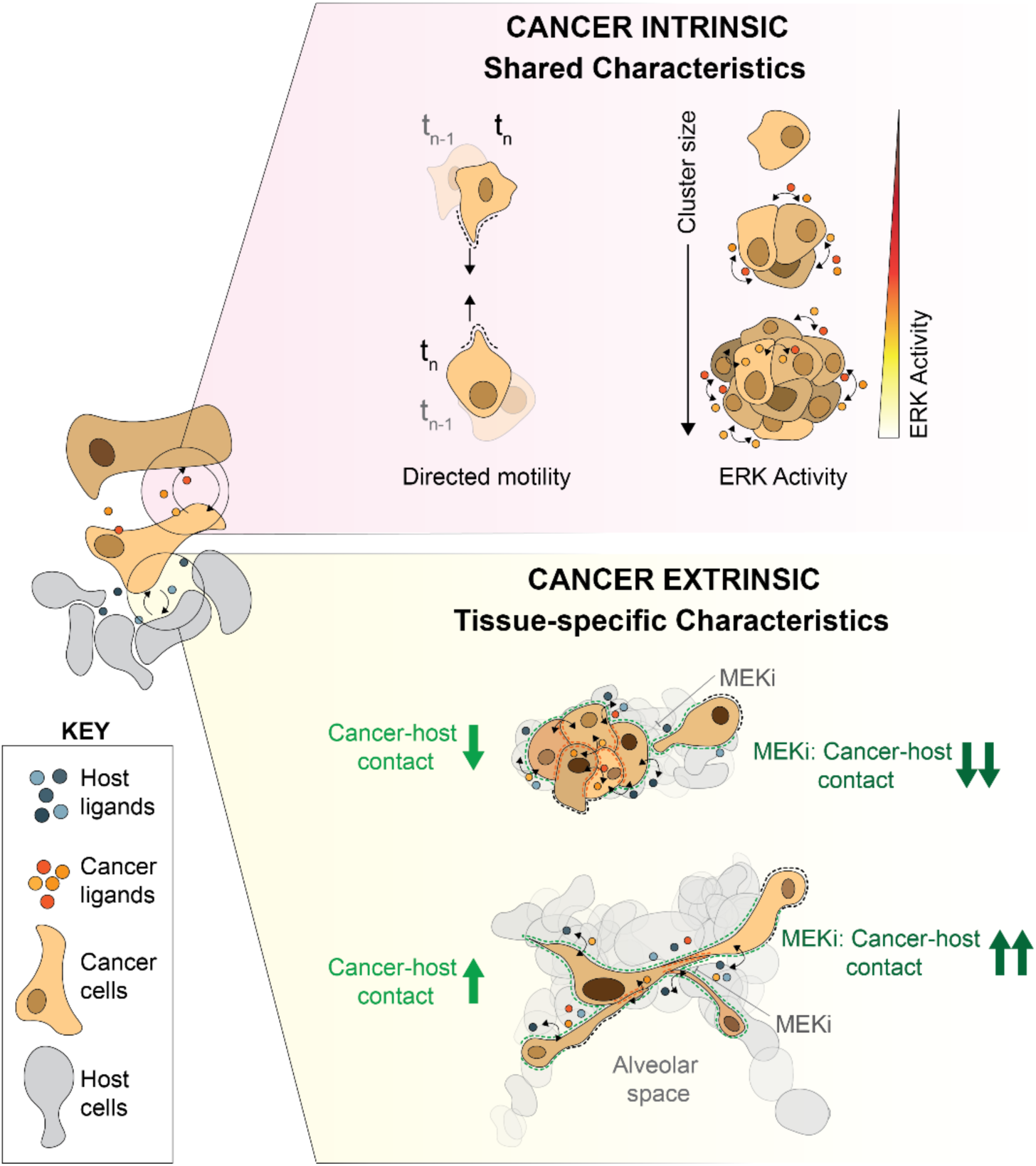
Extrinsic issue context dependent niche formation and ERK-driven plasticity. Schematic summary illustrating converging cancer-cancer and cancer-host paracrine signalling loops driving shared and tissue-specific behaviours in the primary (mammary gland) and metastatic (lung) sites. Shared cancer-cancer paracrine signalling drives cancer cluster formation cluster-size dependent ERK activity, while host environmental context drives differential cancer-host directed cell motility and co-ordinately gives rise to distinct multicellular architectural niches. MEK inhibition differentially modulates cancer cluster morphology and by dampening cancer cell intrinsic properties driven by ERK-mediated ligand secretion, while accentuating the role of tissue context in modulating cancer cell behaviour.

The specific focus of this study was to demonstrate the utility of the novel SITE platform for assessing cell signalling relationships in tissue context. These initial studies provide insight into how microenvironmental interaction context affects the earliest stages of organization and colonization at primary and metastatic locations during breast cancer progression, raising many new questions and opportunities. For example, it remains unclear how signalling activity in the contact niche relates to long-term consequences for cell fate. Linking initial dynamic behaviours observed in the first 24 hours to long-term cell and tumour fate at the extended timescales of disease progression is a key goal moving forward. Because the MammaSITE and LungSITE models can be successfully cultured and imaged for extended periods, this question is answerable, as are questions related to initiation-progression sequences, which are often difficult to study. A key limitation of the current SITE model is that it lacks an immune component. Future iterations will require addition of this important cellular compartment to more faithfully recapitulate the TME *ex vivo*. Nevertheless, this limitation is offset by the capabilities described herein and the intentional modularity of the system itself, as demonstrated with multiple cell lines and models (**Supplementary Figure 1S**). We anticipate that SITE will be easily extended to other cancer-tissue types and adaptable to many biosensors. Future coupling of SITE models with multiplexed imaging and/or dissociative methods, including sequencing and proteomics, promises to yield new and important mechanistic insights by linking live-cell dynamic behaviours, cell-cell interactions, signalling and multimodal molecular readouts in a single platform.

## METHODS

### Cell Culture and Media

HMT-3522 T4-2 cells (obtained from Mina Bissell, Lawrence Berkeley National Laboratory, available on request) were cultured a in DMEM/F-12 (11320033, Gibco) supplemented with hydrocortisone (H4001-1G, Sigma-Aldrich), β-estrogen (E2758-1G, Sigma-Aldrich), sodium selenite (S5261-100G, Sigma-Aldrich), transferrin (T8158-1G, Sigma-Aldrich) and insulin (I1882-100MG, Sigma-Aldrich) as previously described^62^. OS-17 cells^63^ were grown in RPMI (11875093, Gibco) supplemented with 10% foetal-bovine serum (FBS, A4736301, Fisher Scientific). MDA-MB-231 (HTB-26, ATCC) cells were cultured in DMEM supplemented with 10% FBS. All cell lines were maintained at 37°C in 5% CO_2._

### Reporter Line Construction

Reporter T4-2 and MDA-MB-231 cells were generated using lentiviral transduction of ERK translocation reporter (ERK-KTR)^48^, fused to mVenus and selected with puromycin (A1113803, Gibco). OS-17 lines were co-transduced with ERK-KTR fused to iRFP-670, AKT(FOXO)-Neon Green and H2B-mCherry viruses and sequentially selected with puromycin and G418 (10131035, Gibco).

### Lateral tail vein injection

The tail vein injection procedure used was adapted from UCSF (Lateral Tail Vein Injection in Mice and Rats-IACUC Standard Procedure). Briefly, 8-to 12-week-old virgin female NOD SCID IL2R gamma null (NSG, RRID: IMSR_JAX:005557, The Jackson Laboratory) mice were anesthetized with isoflurane (2-3%) until unresponsive, confirmed by the absence of a hind limb withdrawal reflex upon gentle foot pinch. Prior to injection, animals were warmed for 5–10 minutes using a warm water heating pad and/or a heat lamp to facilitate vein dilation. The needle (27-30 gauge) of the syringe containing 100 µL of the injection solution (reporter cells and their corresponding media) was inserted bevel-up into the tail vein, directed toward the head, with the syringe and needle maintained parallel to the tail. The injection was administered slowly, ensuring no resistance was felt. If resistance was encountered or if a blister or white discoloration appeared at the injection site, the needle was withdrawn and reinserted at a location closer to the head. When additional sites were needed, injections were performed on alternate tail veins, progressing from the mid-tail towards the base of the tail. Following injection, the needle was removed, and gentle compression was applied to the site until haemostasis was achieved.

### Mammary intraductal injections (MIND)

MIND injections were performed using an adapted protocol from Kittrell et. al., 2016 under approved protocols by Oregon Health and Sciences University and the Institutional Animal Care and Use Committee^50^. Recipient mice (8-to 12-week-old virgin female NOD SCID IL2R gamma null mice) were purchased from The Jackson Laboratory (Bar Harbor, ME, USA). One to two days prior to the procedure, hair was removed from the abdomen using hair removal cream (Nair^TM^) to expose the inguinal nipples, ensuring no damage to the skin or underlying structures. Mice were anesthetized with isoflurane (2–3%) until unresponsive, confirmed by the absence of a hind limb withdrawal reflex upon gentle foot pinch. The mouse was then placed on a surgical board, and the limbs were secured with surgical tape. The abdominal area was sterilized with 70% ethanol following which the tip (∼0.5mm) of the inguinal nipple was trimmed to allow direct insertion of a Hamilton syringe (1705N, Hamilton). The syringe tip (containing reporter HMT-3522 T4-2 cells or MDA-MB-231 cells and dilute trypan blue (T10282, Invitrogen) for visualization of the injection into the duct) was inserted into the nipple at a 90-degree angle to the animal, and the cell suspension was carefully injected. Following the injection, the mouse was euthanized by cervical dislocation, and thoracotomy was performed to ensure humane euthanasia. The injected mammary glands were collected via dissection and placed into media for further processing. Successful injections were indicated by a thread-like appearance of trypan blue stain throughout the gland and surrounding tissues. In cases of unsuccessful injections, trypan blue stain accumulated only in the mammary fat pad and glands were discarded.

### Generation and maintenance of precision cut lung slices

Mice were anesthetized and euthanized (as described above), following which lungs were harvested by making a vertical incision in the lower abdomen. The diaphragm and rib cage of the mice were removed to allow for complete inflation and perfusion. Mouse lungs were inflated with 2-2.5% low melting point agarose (R0801, Thermo Scientific) dissolved in phenol red-free DMEM/F-12 (21041025, Gibco) with flow rate of about 200µL/second until the lungs were fully inflated. The trachea was held close with a haemostat to prevent backflow of agarose upon perfusion. The perfused lungs were extracted and cooled in ice-cold PBS to allow the agarose to solidify, following which the heart was removed.

Individual lung lobes were cut into 300µm slices with a VF-510-0Z vibratome (Precisionary Instruments). Cut lung slices were placed in advanced-DMEM temporarily. Following this, the PCLS were maintained in defined lung media comprising of DMEM/F-12 supplemented with Hydrocortisone, β-estrogen, sodium selenite, transferrin, insulin, vitamin A (0.2mg/mL, R2625-50MG, Sigma-Aldrich), vitamin C (0.1µg/mL, A4403-100MG, Sigma-Aldrich), primocin (Ant-pm-1, Invitrogen) and Hoechst (1:10,000, 62249, Thermo Scientific).

### Derivation of mammary gland organ fragments

Mammary glands were minced for 30-60 sec with scalpels and digested with Collagenase (C2139-100MG, Sigma-Aldrich), Trypsin (T4799-100G, Sigma), FBS and Y27632 (7230, Stem Cell Technologies) in DMEM/F12 on an orbital shaker for 20 min at 37°C with rotation speed of 170 RPM. The tissue was then manually disrupted and centrifuged for 5 min at 1500 rpm. The supernatant containing fat was separated and further agitated to allow complete digestion of the glands and centrifuged for 5 min at 1500 RPM. The resulting supernatant containing the fat was discarded and the pellet of gland tissue were recombined and suspended in 1% BSA (A7906-50G, Sigma-Aldrich) in DMEM/F12 and centrifuged for 5 min at 1500 RPM to further remove any remaining fat tissue. Finally, the pellet resuspended in a matrix of 70% collagen (354236, Corning) and 30% matrigel (354231, Corning) at a concentration of 1 ductal fragment per microliter of matrix. 20µL organoid-matrix domes were plated in 96-well glass bottom plates (P96-1.5H-N, Cellvis) and allowed to incubate at 37°C with 5% CO2 for 45 min before adding 100 µL of media containing DMEM/F12 supplemented with FGF2 (2.5nM,100-18B-50UG), Primocin, Insulin, β-estrogen (1nM), Hydrocortisone, Transferrin, Y27632 (10µM) per well.

### Live-dead staining and viability assessment of PCLS and mammary organ fragments

To assess viability, PCLS and mammary organ fragments were assayed at 24 hours and 7 days post injection/derivation. Live cells were assessed by Calcein AM L3224, Thermo Fisher Scientific) and MitoSpy (mitochondrial membrane integrity, 424807, Biolegend) staining as described by the manufacturer. A subset of tissue was treated with 70% methanol (322415, Sigma-Aldrich) for 1 hour prior to staining and used as a positive control for dead cells. Tissue survival was determined by measuring the fluorescence intensity of Calcein AM using NIS elements software (Version 5.42.02, Nikon).

### Live-Cell Microscopy

Live-cell imaging for reporter validation was performed on 96-well plates with #1.5 glass bottoms coated using 5 µl of 50 µg/mL rat tail collagen Type I (354236, Corning) dissolved in 20 mM acetic acid LC243501, Cole-Parmer) for one hour then washed with PBS to remove excess collagen. Reporter HMT-3522 T4-2 cells were seeded at a density of 10,000 cells per well and allowed to adhere for one hour prior to adding media and incubated overnight. The following morning, cells were washed twice with phenol-free DMEM/F-12 without additives, then placed in imaging media consisting of phenol-free DMEM/F-12 containing hydrocortisone, β-estrogen, and transferrin, and left to incubate for 4 hours before imaging. Images of each well were collected every 6 minutes for the duration of the experiment. Each image was captured with a Nikon Plan Apo 0.80 NA 20X air objective. Cells were initially imaged for four hours to obtain baseline ERK activity following which they were treated with vehicle (media only), EGF (100ug/mL, AF-100-15-100UG, Thermo Fisher Scientific) or EGFRi (Erlotinib, 2uM, S7786, SelleckChem).

Live-cell microscopy for PCLS and mammary organ fragments was performed on multi-well plates with #1.5 glass bottoms. Mammary organoids were imaged in phenol-free DMEM/F12 supplemented with FGF2, Primocin, B-estrogen, Hydrocortisone, Transferrin, Cellbrite 405 (30105, Fisher Scientific) and Spirochrome 650 (CY-SC501, Cytoskeleton). PCLS were placed in 24-well glass bottom plates (P24-1.5H-N, Cellvis) and clamped flat to the bottom of the plate with custom laser cut acrylic clamps to prevent movement of the tissue during imaging. Lung slices were washed once with sterile PBS following which they were moved into defined PCLS imaging media comprising phenol-free DMEM/F12 supplemented with hydrocortisone, β-estrogen, transferrin, vitamin C, vitamin A, primocin and Hoechst (1:10,000). All plates were imaged on a Nikon Ti-2 inverted microscope fitted with an Oko-Lab environmental chamber maintained at 37°C and 5%CO_2_. Multiple mammary organ fragments or stage positions within lung sections were chosen and time lapse images were captured every 20 minutes with Nikon Plan Apo 40X 0.75 NA objective with a water immersion lens and Fusion-BT camera (Hamamatsu). Both PCLS and mammary organ fragments were treated with vehicle or MEKi (PD0325901, S1036, SelleckChem) for the duration of imaging along with regular media changes with fresh drug. For large format stitched imaging of whole PCLS sections, biopsied PCLS were imaged with the same microscope settings at a singular timepoint. Image capture was automated using NIS-Elements AR software.

### Immunocytochemistry

Reporter HMT-3522 T4-2 cells were plated in glass bottom 96 well plates as described for live-cell preparations. Cells were washed with phenol-free DMEM/F12 with no additives, following which they were placed in imaging media (described above). Cells were treated with media, EGF (100ng/mL) or EGFRi (Erlotinib, 2µM) for 24 hours, following which the treatments were removed and cells rinsed twice with PBS. Cells were fixed in 4% PFA (J19943-K2, Thermo Scientific) for 20 min and permeabilized with 0.1% Triton-X 100 (BP151-500, Fisher Scientific). Cells were blocked with 3%BSA in PBS supplemented with 1% Tween-20 (blocking solution,AAJ20605AP, Fisher Scientific) for 1 hour, following which antibodies against anti-mouse GAPDH (1:200, AM4300, Thermo Fisher Scientific) and anti-rabbit β-catenin (1:200, 8480 Cell Signalling Technologies) were added in blocking solution and allowed to incubate overnight. The following morning, cells were washed with PBS supplemented with 0.5% Tween-20 and secondary antibodies (donkey-anti-mouse-Alexa-647, A-31571, Thermo Fisher Scientific and donkey-anti-rabbit-Alexa-555, A-31572, Thermo Fisher Scientific) were added at a concentration of 1:400 for 2 hours in blocking solution. Cells were then washed with PBS+0.5% Tween-20 and counterstained with Hoechst (1:100) for 10 min. Cells were rinsed with PBS and imaged the Nikon Ti2 automated microscope previously described for live cell imaging. Each sample was scanned at 40x magnification. Identical imaging settings were used for all samples within the same staining.

## COMPUTATIONAL METHODS

### Region of interest extraction

To extract subvolumes for reporter cell and microenvironment analysis, we created coarse masks of reporter cell locations and extracted connected regions of reporter cells over the imaging duration. Coarse segmentation masks were generated from ERK-KTR reporter intensity images using Gaussian smoothing (σ ≈ 13 µm) and intensity-based thresholding (foreground defined as intensities >2.0 standard deviations above background). Segmented regions were filtered by intensity (regions required intensity ≥5.0 standard deviations above background) and size (minimum volume ≈ 6,970 µm³), removing small or dim artifacts. Connected regions across the imaging duration were identified using temporal maximum projections and 3D connectivity analysis, yielding stable reporter cell ROIs. Each identified region was expanded spatially by margins of approximately 6 µm axially and 53 µm laterally to ensure inclusion of surrounding microenvironment context, and the resulting multi-channel, time-resolved subvolumes were exported as individual TIFF stacks for downstream analyses. The total number of datasets, wells and ROIs extracted can be found summarized in **Supplementary Table 1**.

### Reporter cell segmentation

To obtain single-cell cytoplasmic masks, we utilized a deep learning-based segmentation approach using the Cellpose^54^ framework. Representative 2D training slices were generated from three-channel (brightfield, fluorescent reporter, and nuclear stain) 4D imaging stacks from ROIs. Slices exhibiting meaningful intensity variation were selected for manual annotation, yielding a training set containing thousands of annotated images capturing diverse imaging conditions. These annotated datasets were used to train a custom segmentation model initialized from the pre-trained Cellpose cyto3 cytoplasmic model. After training, the optimized model was applied to experimental image stacks, performing full 3D cytoplasmic segmentation. Segmentation parameters included flow smoothing (dP_smooth=2.0) to enhance segmentation robustness, full parameters and cellpose models are available on our provided code repository. Imaging and segmentation data for each ROI was stored within a custom h5 data file for ease of downstream analysis.

### Nuclear segmentation

To quantify nuclear-localization events, such as ERK-KTR translocation, we trained a custom nuclear segmentation model from scratch using Cellpose. Nuclear segmentation training images were generated similarly to cytoplasmic training sets but leveraged distinct imaging information from nuclear fluorescent stains and ERK reporter signals. Thousands of manually annotated 2D training images supported the development of a specialized model trained specifically to identify reporter cell nuclei. Segmentation utilized a nuclear-specific channel combination (fluorescent nuclear stain and ERK-KTR reporter. Segmented nuclear masks were intersected with corresponding cytoplasmic masks to ensure nuclear segmentation specificity within individual cells and stored as binary masks within our h5 file datasets.

### ERK activity quantification

To quantify ERK signalling activity at the single-cell level, nuclear and cytoplasmic intensity ratios of the ERK biosensor fluorescence were computed. Using the previously generated segmentation masks, mean fluorescence intensities within nuclear and cytoplasmic compartments were measured for each segmented cell. The cytoplasmic mask was adjusted to exclude nuclear regions, ensuring disjoint compartments. Background fluorescence was estimated from non-cellular regions in each frame (1st percentile intensity) and subtracted to improve signal accuracy. The ERK nuclear-to-cytoplasmic intensity ratio was calculated per cell to quantify ERK translocation signalling dynamics over time.

### Tissue segmentation

To capture spatial context around individual reporter cells within the local tissue microenvironment, we implemented semi-supervised tissue segmentation using the ilastik random-forest classification framework^43^. Training involved sparse manual labelling of pixels representing three classes—primary tissue (mammary gland or lung), extracellular matrix, and reporter cell regions—using the same image slices employed previously in Cellpose model training. Separate ilastik models were specifically trained for mammary gland and lung tissues to account for their distinct morphological characteristics. Trained ilastik models were applied to full 3D z-stack and time-series imaging data, generating pixel-wise continuous probability predictions for each tissue class. These continuous predictions underwent Gaussian smoothing (σ ≈ 5 pixels) and were thresholded (Δ probability > 0.2 between classes) to produce binary foreground masks, distinguishing reporter cell, primary tissue, and extracellular matrix compartments.

### Boundary Maps

To characterize reporter cell morphology, motility, and interactions with neighbouring cells and the surrounding microenvironment, boundary maps were computed for each segmented cell surface at a resolution of 1.33 µm. Boundaries were extracted from 3D segmentation masks, and geometric features were calculated for each boundary point. After establishing cell lineage connections between consecutive frames, boundary points from matched cells were linked through an optimal transport approach, calculating precise displacements at each boundary point. These displacement vectors enabled quantification of boundary velocities and local surface deformations.

### Cell tracking

To establish temporal trajectories of 3D-segmented reporter cells, we utilized an optimal transport-based tracking method that leverages cell boundary information. At each time point, cell boundaries were extracted, scaled to account for anisotropy between axial and lateral dimensions, and cell centroid positions calculated. Cell identities between consecutive frames were matched by taking neighbouring cells centroid proximity (threshold ≤ 10 µm) with linking the cell in the previous frame with the minimal optimal transport cost between corresponding cell boundaries, computed using the Earth Mover’s Distance (EMD)^55^. This approach explicitly accounts for cell shape changes and local boundary morphology.

### Segmentation and tracking validation

Reporter cell segmentation and tracking validation was performed via manual expert curation. 477 cells were randomly selected spanning imaging duration in each condition. 3D segmentations were evaluated in full in each z-slice and categorized as to whether they matched expert expectations regarding single-cell identification, and overall single-cell boundary morphology. Cell linkages were also manually validated. True positive segmentation and cell identification rates are reported in **Supplementary Table 2**. To quantify false negatives (cells present in ground truth but missed by the model), we manually counted the number of undetected cells across one ROI for each XY position imaged and are reported in **Supplementary Table 3**.

### Contact and microenvironment quantification

Microenvironmental interactions surrounding each reporter cell at every time point were quantified using our established boundary mapping approach. We analysed cell boundary proximity to neighbouring compartments, classifying neighbours into three categories: cancer cells, host tissue cells, and extracellular matrix. For each boundary point, nearest-neighbour distances to each compartment type were computed, and points within 1.9 µm were considered in contact. Note that contact “fractions” are not mutually exclusive between neighbouring boundary types, the same cell boundary point can be within the contact distance cutoff of multiple compartments (e.g. cancer cells and host cells). Cell-level summaries were derived, including the fraction of each cell boundary in contact with each compartment and the average nearest-neighbour distance to each compartment across all boundary points. These spatial metrics—contact fractions and mean distances—were computed for every segmented reporter cell and stored to support subsequent analysis correlating cell signalling and fate decisions to local microenvironmental contexts.

### Microenvironment segmentation validation

To evaluate the accuracy of microenvironmental segmentation generated by the ilastik model, a subset of untrained images from mammary gland organ fragments (13 XY fields, 30 z-slices) were manually annotated using Cellpose3. For lung slices, lung tissue architecture was segmented using Otsu thresholding (4 XY fields, 92 z-slices). The resulting ground truth masks were compared to ilastik model predictions, and segmentation accuracy was quantified by calculating the DICE similarity coefficient, as reported in **Supplementary Table 4**.

### Contact niche states

To identify discrete microenvironmental states based on local cell–cell and cell–tissue interactions, we applied a trajectory-informed kinetic clustering approach to reporter cells characterized by their contact with cancer and host tissue compartments. Each cell at each timepoint was embedded in a 2D contact feature space (cancer and host contact fraction), and fine-grained k-means clustering was performed to define a set of microclusters (n ≈ √N ≈ 220), ensuring sufficient resolution for spatial analysis of signalling and phenotypic trends.

To capture spatial structure in the distribution of dynamic features (e.g., signalling activity, motility), cell-level measurements were aggregated into 20×20 spatial bins using 2D histograms smoothed with a Gaussian kernel (σ = 2 bins). These feature heatmaps were normalized by local cell density to yield interpretable average values across contact niche space.

To model how cells transitioned between microclusters over time, we constructed a Markov state model by assigning each cell at times t and t+1 to its respective microcluster and calculating a transition probability matrix^64^. In this discrete space, a transition matrix T between bins was estimated from the set of transition counts C_ij from microbin i to j as T_ij=C_ij/C_i with C_i=∑_j C_ij. From this transition matrix we computed leading eigenvalues and eigenvectors, and metastable contact niche states were extracted using iterative k-means clustering in this kinetic space^65^. The number of final states was chosen to maximize the sum of the transition timescales representing the metastability of the system. Low-occupancy states (probability < 0.05) were merged into their nearest well-populated neighbour based on Euclidean distance in the kinetic embedding. This approach yielded a set of kinetically distinct, biologically interpretable contact niche states that captured the stable microenvironmental contexts reporter cells occupy over time.

### Centroid motility quantification

To quantify the overall motility of reporter cells, we analysed the displacement of each cell’s centroid over time using cytoplasmic segmentation masks. At each timepoint, the instantaneous displacement vector Δx was computed by subtracting the centroid position at time t from its position at time t - 1, yielding both the magnitude and direction of movement. These displacement vectors were used to calculate motility features including per-frame speed and directional persistence.

To assess how motility is coordinated across space and time, we computed a directional alignment correlation function^66^ between the movement of each cell and its neighbours. Specifically, we evaluated how the displacement vector of a neighbouring cell aligned with the separation vector between the two cells, as a function of spatial distance and time lag. The space-time directional correlation was defined as:

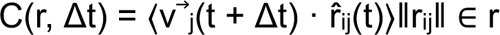

where v⃗ⱼ(t + Δt) is the displacement vector of cell j at time t + Δt, r^ᵢⱼ(t) is the unit vector from cell i to j at time t, and the average is taken over all cell pairs with ǁrᵢⱼǁ within the spatial bin centred at r. Correlations were calculated over radial bins and time lags, with z-axis distances rescaled to account for anisotropic voxel size. The resulting curves quantify how motility alignment decays across space and time, providing a measure of collective migration of cells coming together to form clusters. Heatmaps were smoothed with a Gaussian kernel of 1.0 space and time bins for visual clarity. Unsmoothed versions and a null model with randomly scrambled cell positions are shown in **Supplementary Figure 3C, D.** Error bars in **Supplementary Figure 3B** were calculated by bootstrapping across imaging fields of view and leaving one biological replicate out (𝑛 = 100 per replicate).

### Boundary displacement

To quantify reporter cell motility in terms of localized boundary deformation relative to neighbouring tissue components, we leveraged boundary maps that link individual cell surface points to spatial neighbours and track them over time. For each boundary point of a reporter cell, we calculated the displacement along the direction of the nearest neighbouring surface—either host tissue or extracellular matrix—between successive frames. The boundary displacement in the direction of the neighbour was computed as:

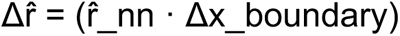

where r^_nn is the unit vector from the reporter cell boundary point to its nearest neighbour surface (host cell or matrix), and Δx_boundary is the boundary displacement vector from time t to t + 1 at that point. Only nearest neighbour surface points directed in the same direction (within 180 degrees) of the outward facing surface normal were included in the calculation.

Displacements were binned according to boundary separation distance (0–40 µm, 15 bins) and plotted as mean values of the signed magnitude of boundary motion—positive values indicating extension toward the neighbouring structure, and negative values indicating retraction. Single-cell distributions span negative and positive values, all positive average displacements are consistent with a small net increase in cancer-cancer and cancer-host contacts over the experiment duration, and also a near steady-state balance of directed cell-cell attraction and stochastic motility. Outlier displacements larger than 40 µm were discarded. Error bars in **Figure 4D-E** were calculated by bootstrapping across imaging fields of view and leaving one biological replicate out (𝑛 = 100 per replicate).

### Cluster size description

To characterize the size of reporter cell clusters, connected coarse connected regions in the ERK-KTR intensity volumes were extracted. At each timepoint, the ERK-KTR intensity channel was first smoothed using a 3D Gaussian filter (σ = 10 µm in the lateral dimension, adjusted by z-scale axially), and a z-score–normalized foreground mask was generated using a threshold of z = 2.0. Connected components within this foreground mask were labelled to define multicellular clusters. Within each frame, labelled cytoplasmic segmentations were then assigned to ERK-KTR-defined clusters by intersecting reporter cell labels with each connected region. Cells belonging to the same labelled component were grouped under a common cluster ID. The number of cells assigned to each cluster was used as the cluster size, and this information was stored for all reporter cells across frames.

### ERK activity spatiotemporal quantification

To analyse the dependence of ERK activity triggered ERK signalling, we characterized spatiotemporal correlations in ERK signalling activity. A discrete Markov model was constructed using paired single-cell ERK-KTR C/N values from consecutive timepoints and binned into n = 41 equally spaced intervals between 0 and 3. The eigenstructure of the resulting transition matrix revealed slow, metastable dynamics and was used to identify robust boundaries between “low” and “high” ERK activity. Based on the extrema of the dominant eigenvectors, we defined C/N thresholds at 0.75 and 1.55, with a midpoint of 1.15 used to binarize single-cell ERK states.

To quantify spatiotemporal organization in ERK activity, we computed a radial correlation function^66^ focused on the spatial propagation of activity from ERK “on” cells. For each pair of cells i and j, with Sᵢ(t) ∈ {0,1} indicating ERK activity at time t, we computed:

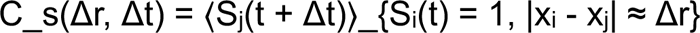

This represents the average probability of a cell being ERK “on” at a spatial distance Δr and temporal lag Δt, conditional on being near a cell that was “on” at time 0. Only cells that were ERK-active at the initial timepoint contributed to the signal source set (i.e., Sᵢ(t) = 1), allowing the interpretation of Cₛ as a conditional expectation of ERK propagation. Correlations were measured in n = 20 radial bins up to a maximum interaction distance of ∼75 µm, and across time lags Δt = 0–27 frames. Heatmaps were smoothed with a Gaussian kernel of 1.0 space and time bins for visual clarity. Unsmoothed versions and a null model with randomly scrambled cell positions are shown in **Supplementary Figure 4A.** Error bars in **Supplementary Figure 4B** were calculated by bootstrapping across imaging fields of view and leaving one biological replicate out (𝑛 = 100 per replicate).

To assess how ERK activity depends on local cell density, we also stratified cells by ERK-KTR cluster size, defined by connected components of binarized ERK-KTR activity in 3D volumes. For each size bin (0–3+ cells), we calculated the fraction of cells that were ERK-active. The ERK “on” fraction increased monotonically with cluster size, indicating that local density and spatial context shape the likelihood of ERK activation in tissue.

### ERK activity dependence upon local signalling S/R

To capture the dependence of ERK signalling on the local reporter cell signalling density, we treated ERK-on reporter cells as sources of soluble ligands. On spatiotemporal scales shorter than the estimated diffusion length (∼110 μm), the ligand profile from a continuous point source approximates a steady-state distribution, decaying radially as ∼ 1⁄r. This assumption is supported by the diffusion coefficient for typical signalling molecules (𝐷 ≈ 10 μm²/s) and a frame interval of 20 minutes, yielding a characteristic diffusion length:

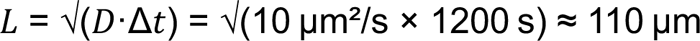

Based on this, the local signalling environment surrounding a given reporter cell was quantified by summing the 1⁄r-weighted contributions of all neighbouring ERK-on cells within a radius 𝑟ₘₐₓ. This effective signal score, denoted 𝑆/R, represents the concentration-like contribution of ERK-on neighbours and is defined as:

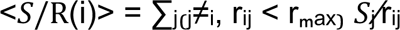

where 𝑆ⱼ = 1 if the neighbour cell is ERK-on and 0 otherwise, and rᵢⱼ is the spatial distance between cells i and j, normalized by the average cell diameter R. Cells were classified as ERK-on or ERK-off using a threshold on the ERK cytoplasm-to-nucleus (c/n) ratio, set at ERKT c/n=1.15. The spatial signal 𝑆/R was then compared to binarized ERK activation across the population to assess whether ERK activity was enhanced in locally signalling-dense neighbourhoods.

### Spatiotemporal Markov model of ERK signalling

To characterize the cluster size dependent ERK signalling activity relationships in the lung and gland, we modelled ERK activity with spatiotemporal resolution via a dynamical model of ERK on and off kinetics, coupled to local ERK signalling density. Reporter cells were classified into discrete kinetic states using a two-dimensional state space defined by local S/R (signal-to-radius) contributions and binarized ERK activity, with the latter thresholded at cytoplasm-to-nucleus (c/n) ratio > 1.15. Reporter cells were first grouped by local S/R using k-means clustering (𝑘 = 5), with centres spaced across the observed S/R range. Each S/R cluster was then paired with both ERK-on and ERK-off states to define a total of 2𝑘 = 10 microstates in the Markov model.

Cell trajectories (𝑥₀, 𝑥₁) were assigned to these microstates across consecutive timepoints, and transition matrices 𝑃 were estimated for each tissue type. Probabilities for ERK activation (𝑃_on_ = 𝑃(state_off → state_on)) and deactivation (𝑃_off_ = 𝑃(state_on → state_off)) were computed from the counts of these events as described above for Markov model construction. Bootstrapping across imaging fields of view and leaving one biological replicate out (𝑛 = 100 per replicate) provided 95% confidence intervals for transition probabilities.

To test whether the observed ERK activation levels were consistent with a spatially coupled ligand signalling model, we simulated randomized populations of cells seeded via Poisson-disk sampling on a sphere. Cells were initialized as ERK-off at-24 hours, and ERK states were iteratively updated using the fitted Markov model and S/R scores computed at each step. ERK active fractions were taken as averages over t=0-24 hours. Over a range of cell densities, the simulated ERK activation fraction increased with cluster size similar in scale and variance with experimental data, confirming that S/R signalling can account for density-dependent ERK activation behaviour, supporting the view that local ERK signalling density reinforces sustained ERK activation.

## Code Availability

The custom code for image processing and analysis can be found via GitHub at (https://github.com/cancer-dynamics/SITE_Murthy_2026) sample subset of the data and models used for image processing and analysis are available via kaggle at (https://doi.org/10.34740/kaggle/dsv/13784905)

## ACKNOWLEDGEMENTS

This project was supported by funding (Grant No. 2023-1713) from the Cancer Early Detection Advanced Research Center at Oregon Health & Science University’s Knight Cancer Institute (Jeremy Copperman and Alexander Davies). Alexander Davies acknowledges support from National Institutes of Health, Office of Research Infrastructure Programs, under award number K01OD031811. The research reported in this publication used computational infrastructure supported by the Office of Research Infrastructure Programs, Office of the Director, of the National Institutes of Health under Award Number S10OD034224. The content is solely the responsibility of the authors and does not necessarily represent the official views of the National Institutes of Health. We would like to thank Jason Ware for assistance with the design and preparation of acrylic clamps and Laura M. Heiser, Daniel M. Zuckerman and Ellen Langer for their thoughtful discussions and helpful suggestions on the manuscript.

---

**Supplementary Figure 1:**
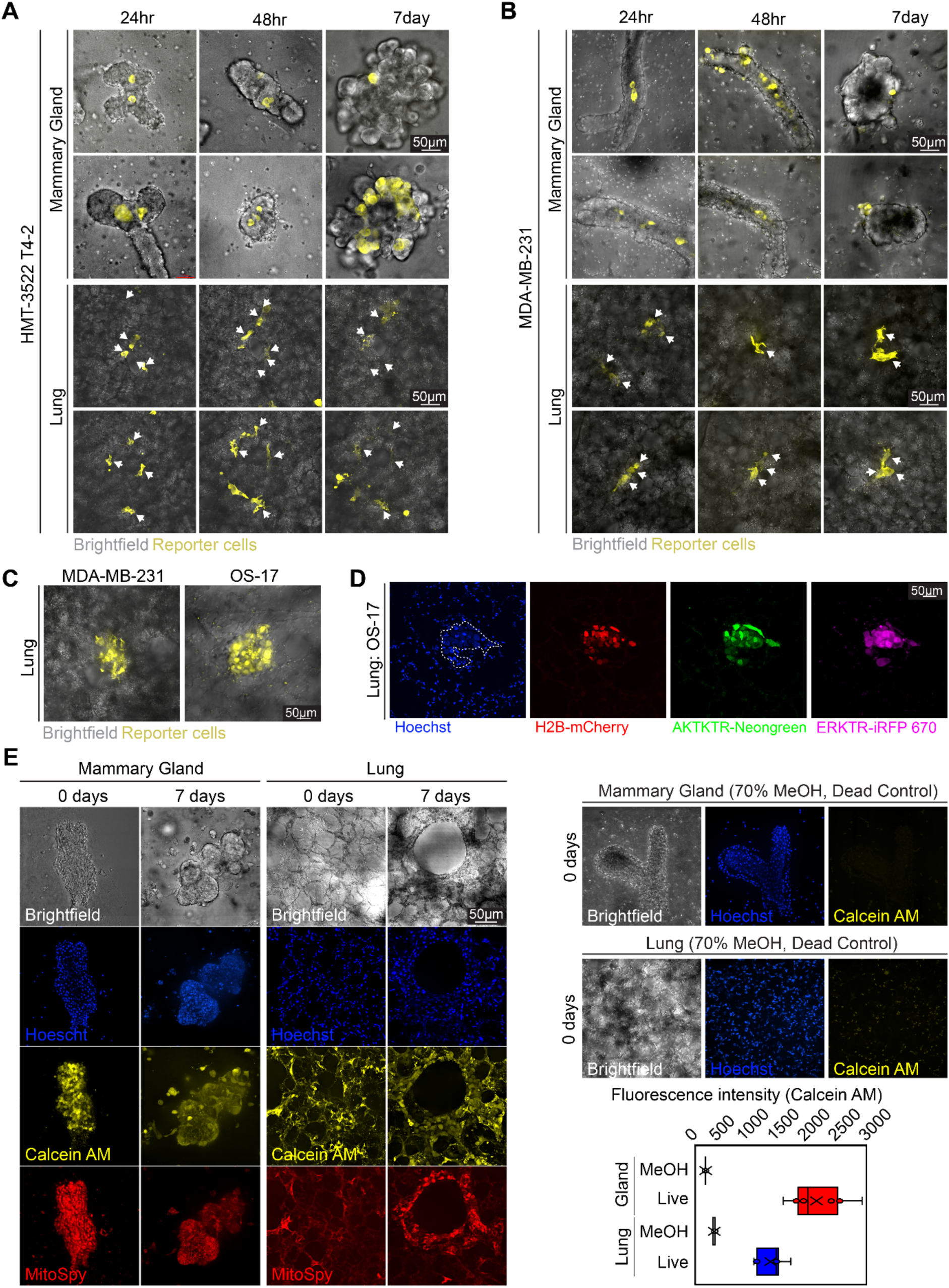
(A) Representative maximum projection intensity images of brightfield (grey) and fluorescence overlay images showing engraftment and distribution of reporter T4-2 cells (yellow) in *ex vivo* mammary organ fragments (top) and PCLS (bottom) captured at indicated timepoints (24hr, 48hr or 7days). Reporter cells in lung images are represented by white arrows. **(B)** Representative maximum projection intensity images of brightfield (grey) and fluorescence overlay images showing engraftment and distribution of reporter MDA-MB-231 cells (yellow) in *ex vivo* mammary organ fragments (top) and PCLS (bottom) captured at indicated timepoints (24hr, 48hr or 7days). Reporter cells in lung images are represented by white arrows. **(C)** Representative maximum projection intensity images of brightfield (grey) and fluorescence overlay images showing engraftment of MDA-MB-231 (left) and OS-17 cells (right) following tail-vein injection and harvest of lungs two weeks post-injection. **(D)** Representative maximum projection intensity images of OS-17 tumour cells with H2B-mCherry, AKT-KTR Neongreen and ERK-iRFP-670 reporters following tail-vein injection and harvest of lungs 2 weeks post-injection. **(E)** Representative maximum intensity projection images of Calcein AM (yellow) and MitoSpy (mitochondrial activity, red) 24 hours (0 days) and 7 days post generation of mammary organ fragments and precision cut lung slices (left). Negative control mammary organ fragments and PCLS were treated with 70% methanol for 1hour at room temperature (right). Calcein AM fluorescence intensity was quantified from three independent experiments (N=3).

**Supplementary Figure 2:**
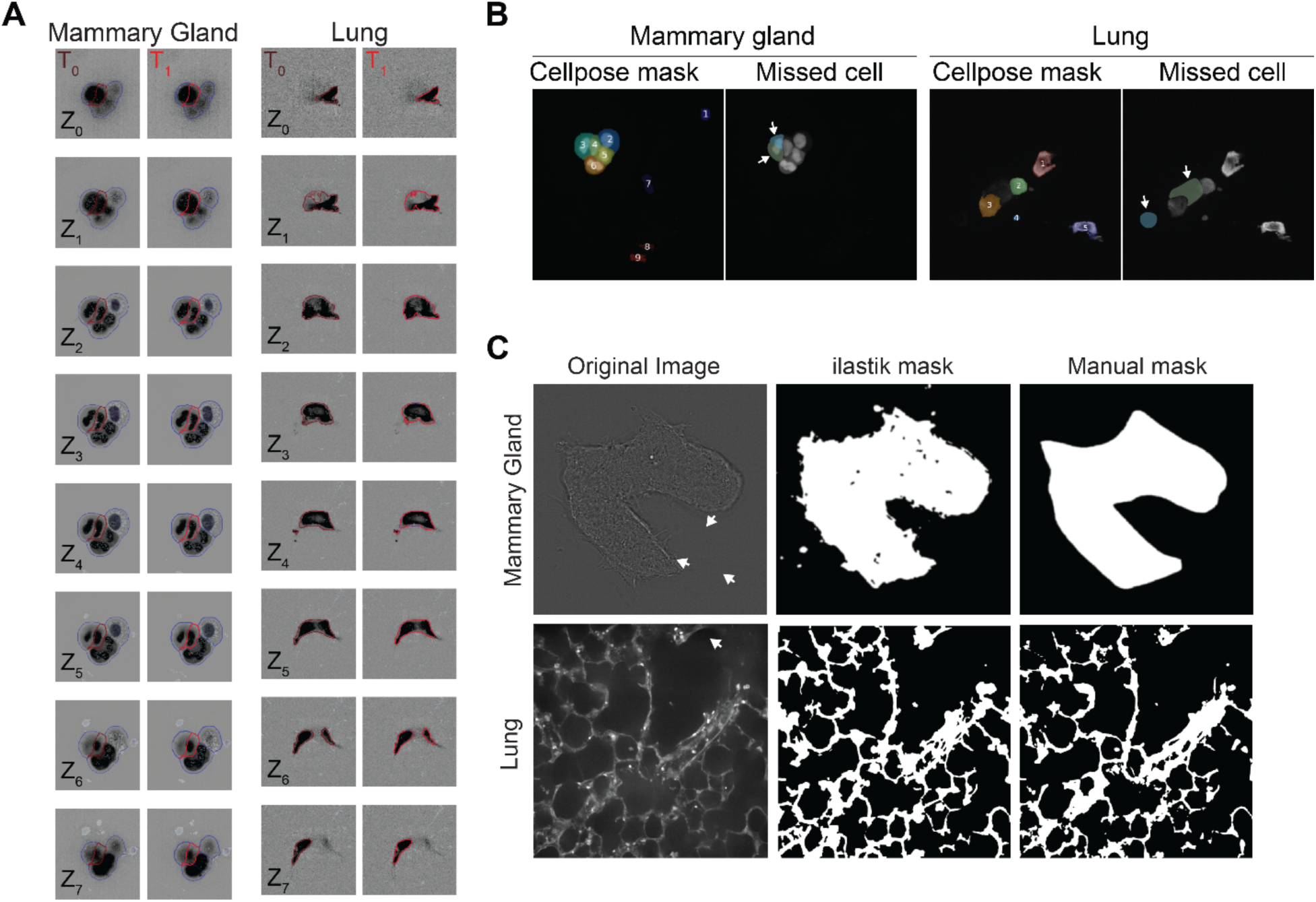
(**A**) Representative 3-D segmentation tracking of single cells in each successive axial z-slice between T_0_ and T_1_ for both mammary gland organ fragments and the PCLS. Blue outlines denote all segmented cells in the field of view, while the red outline highlights the specific cell being tracked across time. (**B**) Validation of Cellpose-based cytoplasmic false negative segmentation of cancer cells in mammary gland organ fragments and PCLS. Representative images comparing automated segmentation results from Cellpose3 to manually annotated ground truth missed masks. White arrows indicate incorrect/missed segmentations from trained Cellpose3 model. (**C**) Validation of ilastik-based segmentation in mammary gland and lung tissues. Representative examples of segmentation performance for the trained ilastik model in mammary gland and lung models. Ilastik segmentations for both models (middle) were generated from raw images (left) and validated against masks made via manual (right) annotation (mammary gland) or via thresholding (lung). White arrows indicate over-segmented areas in the ilastik model.

**Supplementary Figure 3:**
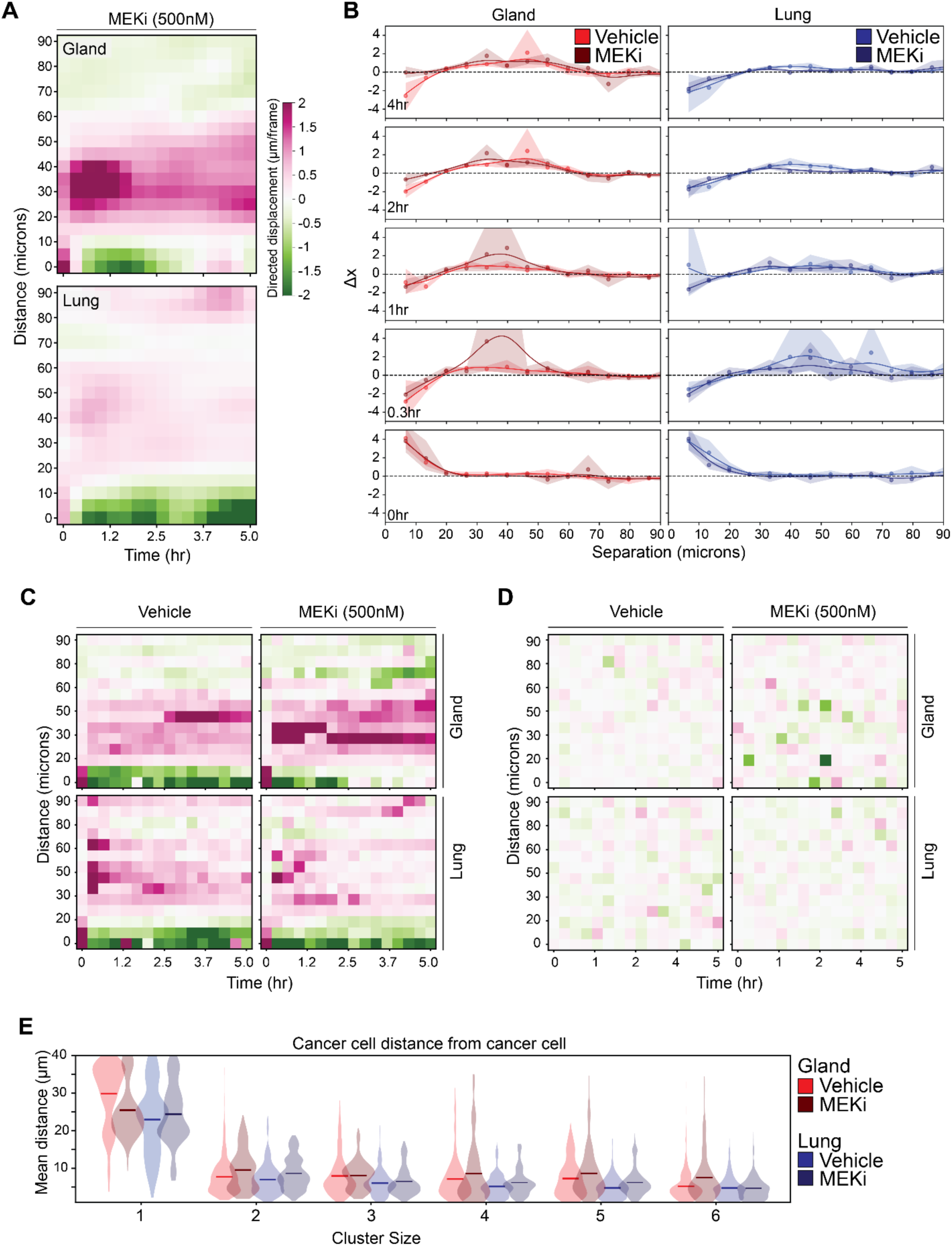
(**A**) Heatmaps illustrating the time evolution of on-axis displacement (μm/frame) as a function of cell-cell distance (microns) over 5 hours in the mammary organ fragments (top) and PCLS (bottom) under MEKi (PD0325901, 500nM). (**B**) Plots of mean on-axis displacement (Δx) versus separation distance (microns) at indicated time points (0, 0.3, 1, 2, 4, 8 hours) for mammary organ fragments (red) and PCLS (blue) microenvironments under vehicle control (light red, light blue) and MEKi (dark red, dark blue) conditions. Shaded regions represent the standard error of the mean. (**C**) Non-smoothed heatmaps of on-axis displacement (μm/frame) as a function of cell-cell distance (microns) over 5 hours in the mammary organ fragments (top) and PCLS (bottom) under vehicle and MEKi conditions. **(D)** Null heatmaps of on-axis displacement (μm/frame) as a function of cell-cell distance (microns) over 5 hours in the mammary organ fragments (top) and PCLS (bottom) under vehicle and MEKi conditions. (**E**) Violin plots of cancer–cancer centroid distances, stratified by cluster size in mammary organ fragments (red) and PCLS (blue) in vehicle (light red, light blue) and MEK inhibitor treated (dark red, dark blue) conditions. Solid bars represent mean of all plotted cells within the cluster size classification (see Methods). Significance was determined by means of Welch’s t-test (***-P≤0.0001, **-P≤0.001) on individual ROIs. A summary of pairwise statistics can be found in Table 5.

**Supplementary Figure 4:**
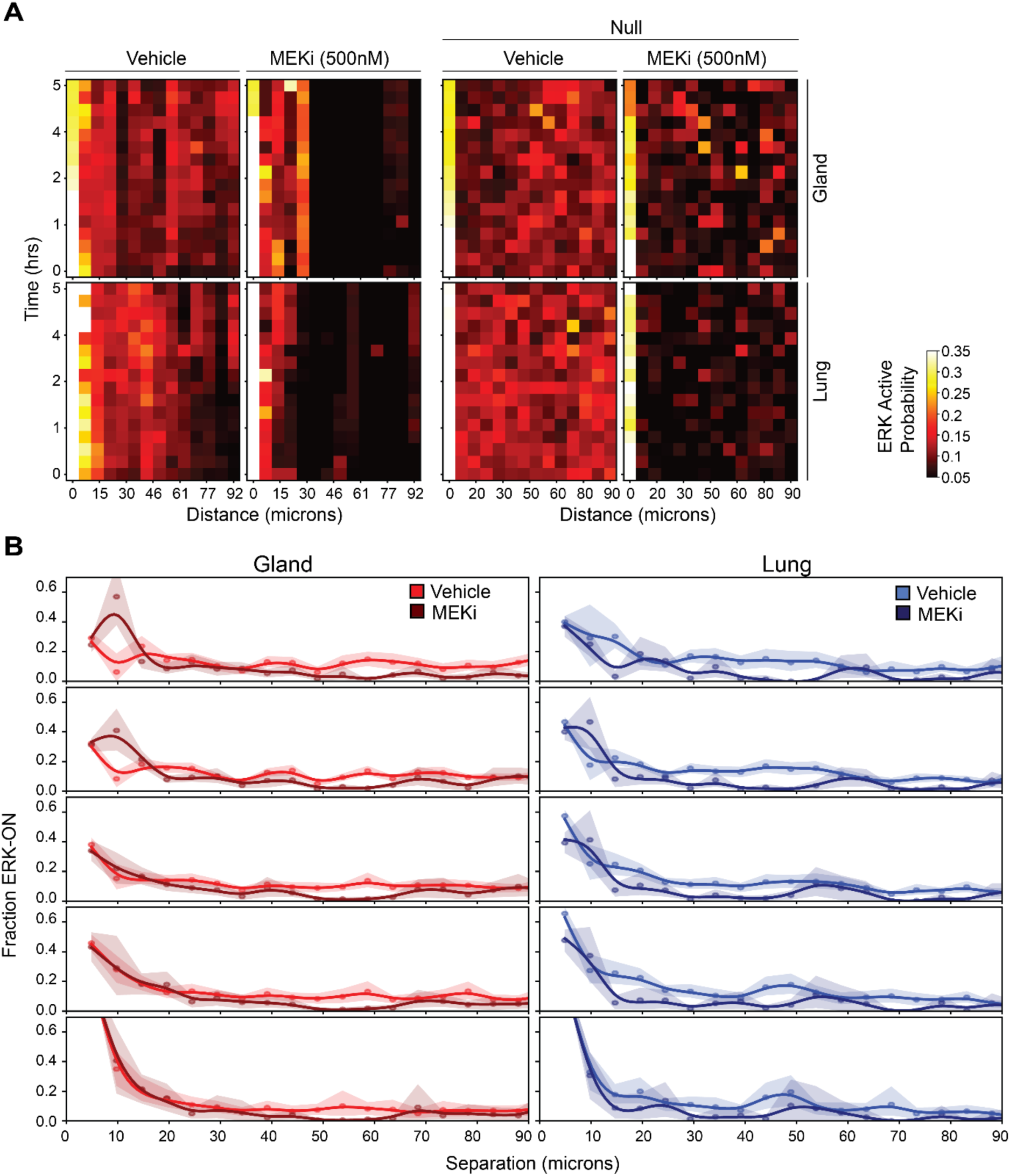
(A) Heatmaps of spatiotemporal ERK activation in vehicle vs. MEK inhibitor (PD0325901, 500nM) treated conditions (left) vs null heatmaps (right). Graphs represent ERK activation probability as a function of time and distance from neighbouring ERK_ON_ cell. **(B)** Line plots show the fraction of ERK-ON cancer cells as a function of separation distance (microns) from the ERK_ON_ cell neighbour, measured at five time points (0, 0.3, 1, 2, and 4 hours) in the mammary organ fragments (left, red) and PCLS (right, blue) microenvironments in vehicle-treated (lighter lines) and MEK inhibitor (PD0325901, 500 nM, darker lines) conditions. Graphs represented as mean and 95% CI (shaded region).

**Supplementary Figure 5:**
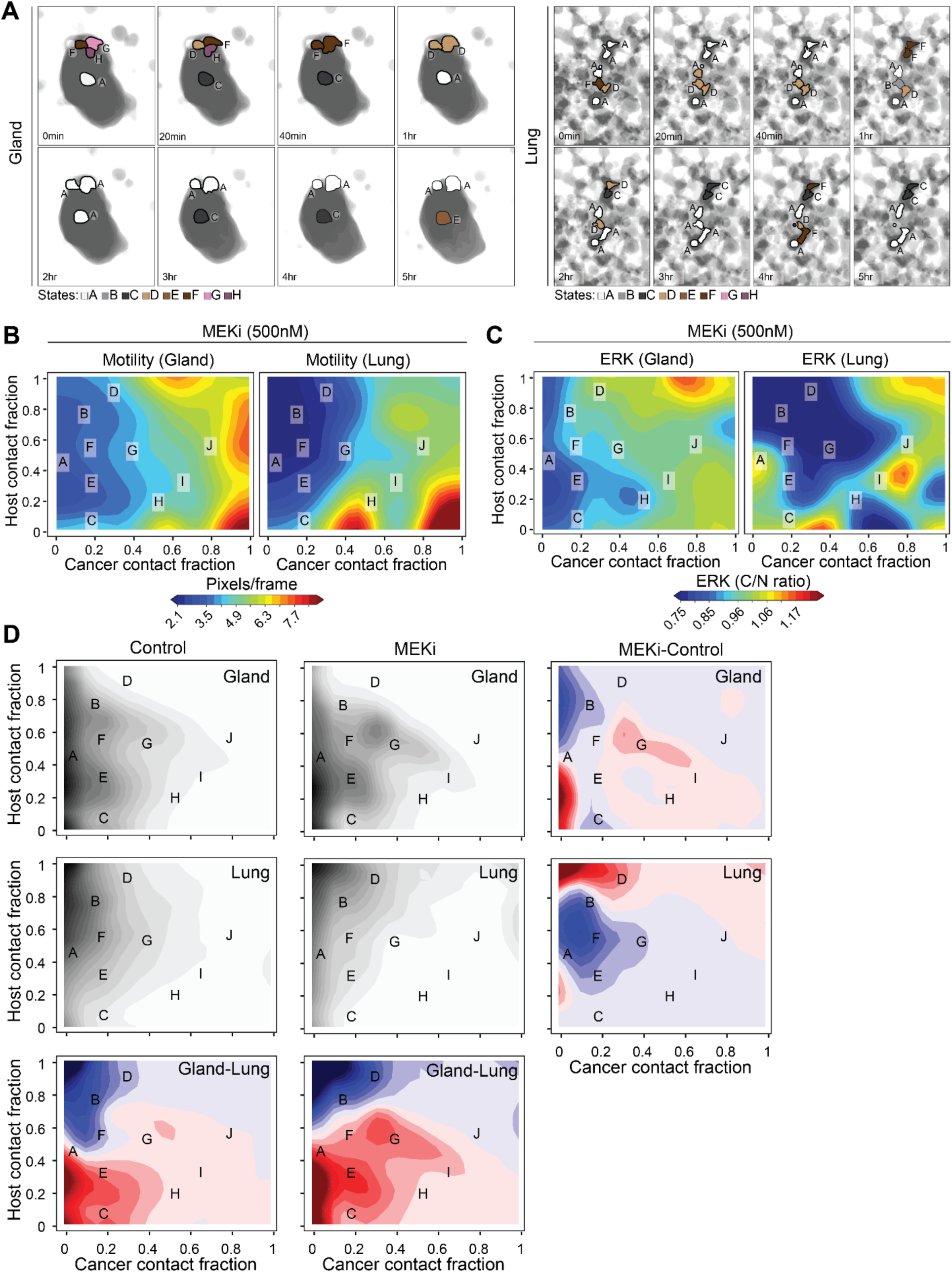
(A) Representative time lapse of maximum intensity projection of segmented cell and microenvironment masks color-coded by assigned contact niche state (states A-H) in both mammary organ fragments (left) and PCLS (right). **(B)** Heatmaps depicting motility (pixels/frame) of cancer cells in the mammary organ fragments (left) and PCLS (right) under MEK inhibitor (PD0325901, 500 nM) treatment, plotted as a function of cancer contact fraction (x-axis) and host contact fraction (y-axis). Distinct contact niche states (A–J) are labelled within the contact space. **(C)** Corresponding heatmaps showing ERK activity (cytoplasmic-to-nuclear ratio) in the mammary organ fragments (left) and PCLS (right) under MEK inhibitor (PD0325901, 500 nM) treatment, mapped across the same contact niche space as in (B). **(D)** Density contour plots depicting the distribution of contact niche states across the mammary gland and lung models under control and MEK inhibitor (MEKi) conditions. Difference maps (right) illustrate the effect of MEKi, (bottom) and gland-lung environmental context.

**Supplementary Table 1:** Summary of imaging datasets used for analysis. Number of XY positions and ROls analyzed per mouse (left) and total XY counts by treatment condition (right) for MIND and tail-vein injections.

|  | nXYs | nROIs |  | Vehicle | MEKi |
| --- | --- | --- | --- | --- | --- |
| **MIND Injection** |  |  | MIND Injection | nXY= 32 | nXY= 19 |
| 1 | 7 | 15 | Tail-vein Injection | nXY= 14 | nXY= 14 |
| 2 | 11 | 12 |  |  |  |
| 3 | 13 | 29 |  |  |  |
| 4 | 11 | 17 |  |  |  |
| 5 | 9 | 14 |  |  |  |
| **Tail-vein Injection** |  |  |  |  |  |
| 1 | 10 | 32 |  |  |  |
| 2 | 8 | 16 |  |  |  |
| 3 | 10 | 22 |  |  |  |

**Supplementary Table 2:** Evaluation of segmentation and tracking performance in ex vivo mammary gland and lung microenvironments. Segmentation accuracy was determined by comparison of automated cell and tissue segmentation results to manually annotated ground truth images. Tracking accuracy was assessed by the percentage of correctly linked cell tracks over time. Values are reported as percentages for each tissue type.

|  | Mammary Gland | Lung |
| --- | --- | --- |
| Segmentation Accuracy | 89.6% | 85.7% |
| Tracking Accuracy | 97.0% | 97.5% |

**Supplementary Table 3:**
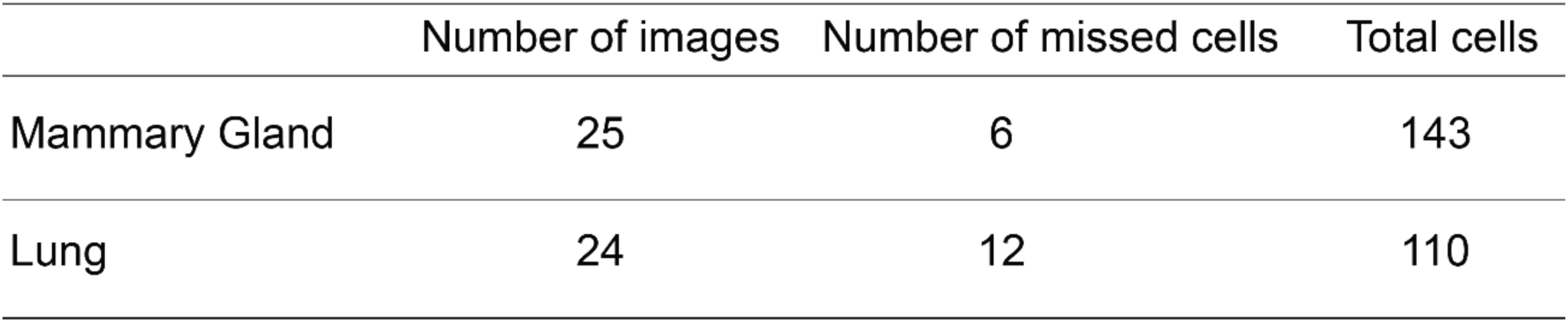
Evaluation of cellpose3 segmentation model.

|  | Number of images | Number of missed cells | Total cells |
| --- | --- | --- | --- |
| Mammary Gland | 25 | 6 | 143 |
| Lung | 24 | 12 | 110 |

**Supplementary Table 4:** Evaluation of iLASTIK model of segmentation in gland and lung microenvironments.

|  | Number of XYs | Number of z-slices | DICE score |
| --- | --- | --- | --- |
| Mammary Gland | 13 | 30 | 0.86±0.04 |
| Lung | 4 | 92 | 0.70±0.03 |

